# A covalent anti-HIV compound induces HIV-1 capsid multimerization and degradation, decomposing the viral core

**DOI:** 10.1101/2025.04.19.649630

**Authors:** Tomofumi Nakamura, Nobutoki Takamune, Mayu Okumura, Jun-ichirou Yasunaga, Masaharu Sugiura, Masayuki Amano

## Abstract

Covalent inhibitors form strong, essentially irreversible interactions with their targets, offering enhanced structural selectivity within the target cavity and prolonged target engagement even at low doses. Here, we report that ACAi-001, an HIV-1 capsid (CA)-targeting compound identified through *in silico* screening of the hydrophobic cavity in the CA N-terminal domain (CA-NTD), inhibits HIV-1 replication in cell-based assays. ACAi-001 covalently binds CA to induce aberrant CA multimerization and degradation *in vitro*, a pharmacological property distinct from that of existing CA inhibitors such as lenacapavir. This mechanism was confirmed by Western blotting (WB), size exclusion chromatography (SEC), differential scanning fluorimetry (DSF), and liquid chromatography-mass spectrometry (LC-MS). These results suggest that ACAi-001, with two putative covalent interactions in its chemical structure, binds directly to Ser16 in the N-terminal domain targeting cavity in addition to Cys198 or Cys218 in the C-terminal domain of CA. The treatment of viral particles with ACAi-001 results in the loss of intact, conical HIV-1 cores, as directly visualized by transmission electron microscopy (TEM), establishing the HIV-1 core as the direct target of ACAi-001. These findings demonstrated that ACAi-001 may be a promising next-generation anti-HIV-1 CA inhibitor with unique mechanisms of action, offering new insight into HIV-1 core assembly and disassembly.

**Graphical abstract:** 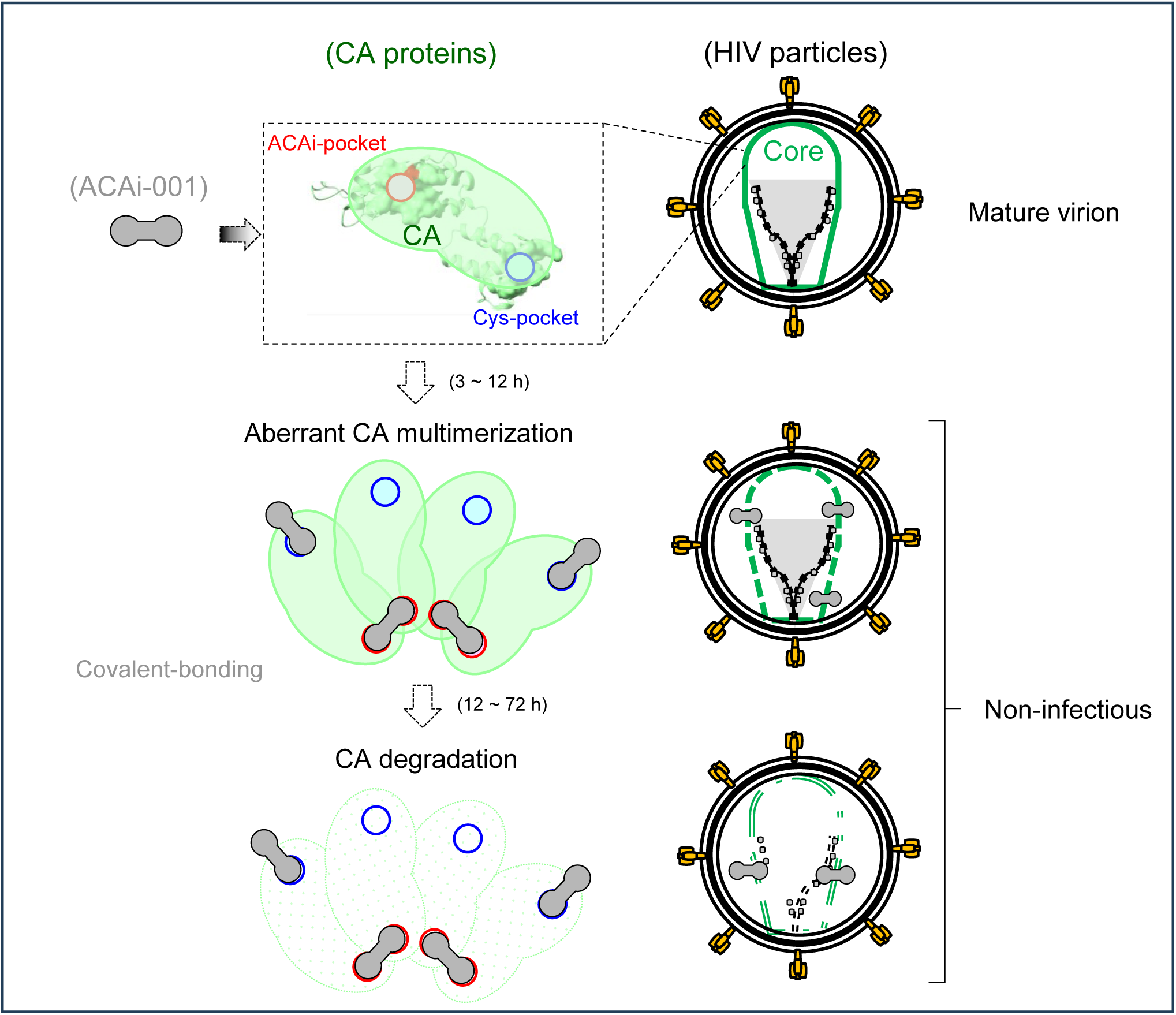

---

During human immunodeficiency virus type 1 (HIV-1) replication, immature virions bud from infected cells and begin to mature through the proteolytic processing of Gag and Gag-pol precursor proteins. HIV-1 protease (PR) cleaves the Gag precursor Pr55 into three distinct structural proteins [matrix (MA, p17), capsid (CA, p24), and nucleocapsid (NC)] and three peptides [spacer peptides 2 and 1 (p2 and p1, respectively), and p6] (1, 2). CA, cleaved from the Gag polyprotein, assembles into hexamers and pentamers that constitute the mature cone-shaped CA (HIV-1 core) (3, 4). We previously reported that amino acid (AA) insertions at specific positions in HIV-1 CA cause significant CA degradation and that HIV-1 variants carrying such AA insertions significantly impaired infectivity and replication capacity. We hypothesized that the observed CA degradation was due to conformational and structural incompatibility caused by AA insertion at a specific site on CA (5). Small molecules can interact with target proteins through various binding modes; recently, potent interactions such as covalent binding have attracted attention (6). Covalent drugs react covalently with their target proteins and irreversibly inhibit their biological functions. Drugs such as aspirin and penicillin are well-known examples of covalent drugs (7) that were not intentionally designed as such covalent drugs. Covalent drugs exert strong and sustained effects by irreversibly binding to their target proteins. In addition, they can react with specific amino acid residues in proteins to selectively inhibit protein families or subtypes, even though non-covalent small-molecule ligands interact poorly with them (8). In contrast, covalent drugs may cause side effects by reacting nonspecifically with off-target proteins or other biological components (9). In the last decade, new types of covalent drugs have been developed that are precisely and molecularly designed to react with target proteins, known as targeted covalent inhibitors, for anticancer therapy (7, 10).

Based on these observations, we predicted that a compound tightly bound to a specific degradation-inducing region(s) of CA could lead to CA degradation and suppress viral replicability. In the present study, we screened a library of millions of commercially available compounds via *in silico* docking simulation for binding to such hydrophobic regions on the CA surface, evaluated the anti-HIV-1 activity of compounds with favorable docking scores, and investigated their effects on CA proteins expressed in *Escherichia coli*, 293T, or COS7 cells, as well as on viral lysates of HIV-1_NL4-3_. Here, we characterize the unique *in vitro* pharmacological properties of the resulting lead compound, ACAi-001, which covalently targets the CA-NTD hydrophobic cavity and induces aberrant CA multimerization and degradation, and discuss the implications of this mechanism for HIV-1 CA-targeted drug development. As a chemical tool, ACAi-001 may also help probe the structural requirements of HIV-1 CA stability and core integrity and reveal the functional role of a CA cavity not previously targeted during HIV-1 replication.

## Results

### ACAi-001 binding profile to the hydrophobic target cavity of CA-NTD

As we previously reported, an AA insertion between Arg18-Thr19 near the target cavity of HIV-1 CA causes conformational and structural incompatibility of CA, leading to CA degradation (5). We hypothesized that a small compound that binds tightly or covalently to the target cavity could induce CA degradation. The hydrophobic cavity at the N-terminal domain of CA (CA-NTD) had a suitable size (252.8 Å^3^) based on crystallographic data of the CA monomer (Figure 1A) (11) and had sufficient space for certain small compounds to potentially fit snugly in the cavity (Figure 1B). This target cavity of the CA-NTD in the center of the CA hexamer may also affect CA hexamer formation (Figure 1C). We then selected 6,842,684 compounds from a virtual library of 8,555,483 commercially available compounds based on parameters predictive of favorable oral pharmacokinetics, including the number of H-bond donors and acceptors, molecular weight, calculated log P, and number of rotatable bonds, criteria related to absorption, distribution, metabolism, and excretion (ADME) (Table 1). Next, the simulated binding scores of each compound to the target cavity were calculated using a virtual docking simulation algorithm, and compounds with good binding scores were further evaluated using cell-based HIV-1 drug susceptibility assays (Figure S1A). To date, we have identified more than 40 active compounds against HIV-1_LAI_ (12) using a 3-(4,5-dimethylthiazol-2-yl)-2,5-diphenyltetrazolium bromide (MTT) screening assay; these compounds showed no or low cytotoxicity against MT-2 and MT-4 cells which are lymphoblastic leukemia-derived T cells. One of these compounds, ACAi-028, binds noncovalently to the target cavity (13) and exerts anti-HIV-1 activity. The chemical structure of ACAi-001 and top 10 putative binding poses of ACAi-001 to CA are shown in Figure 1A. In this study, ACAi-001, which exhibited potent anti-HIV-1 activity (Table 2) and lower cytotoxicity (Table 3), was investigated to understand the details of its anti-HIV-1 activity.

**Figure 1.**
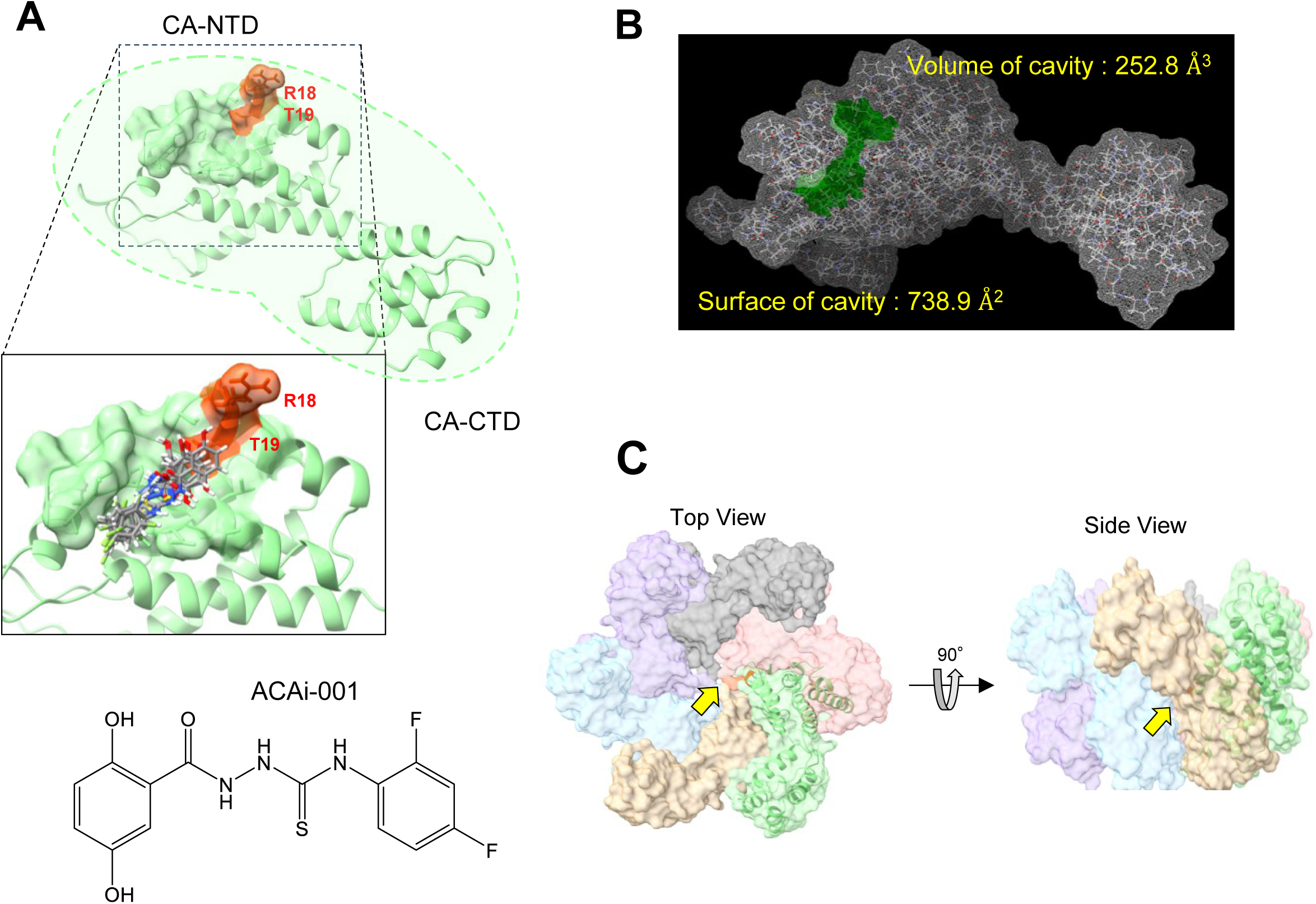
ACAi-001 was identified by targeting a hydrophobic cavity near Arg18-Thr19 on the surface of the CA N-terminal domain (NTD). (**A**) The CA monomer comprises the CA-NTD and CA-CTD; the Arg18–Thr19 region is highlighted in red. Ten representative docking poses of ACAi-001 in the CA-NTD hydrophobic cavity (PDB: 4XFX) and its chemical structure are shown. (**B**) Surface (Å^2^) and volume (Å^3^) of the targeted cavity. (**C**) Location of the target cavity (yellow arrow) within the CA hexamer (PDB: 8CKV), shown from top and side views. Docking simulations were performed using SeeSAR and FlexX v14.2.1 (BioSolveIT GmbH, Sankt Augustin, Germany).

**Table 1.** Factors and characteristics of drugs in oral bioavailability prediction.

| Drugs | MW (Da) | LogP | CLogP | LogS | TPSA ($\text{\AA}^2$) | pKa |
| --- | --- | --- | --- | --- | --- | --- |
| ACAi-001 | 339.32 | 2.59 | 0.61 | -4.75 | 93.62 | 7.72 |
| PF74 | 453.59 | 4.39 | 5.18 | -6.42 | 43.86 | N/A |
| LEN | 968.28 | 5.61 | 3.47 | -12.72 | 152.97 | N/A |

\*Topological Polar Surface Area, TPSA
LogP (octanol/water partition coefficient) indicates fat/water solubility; CLogP is its calculated value, with higher values indicating greater fat solubility. LogS indicates water solubility, with higher values indicating greater solubility. TPSA approximates a molecule's polarized surface area; larger TPSA values indicate lower membrane permeability. pKa, acid dissociation constant. Values were calculated using ChemDraw ver. 17 (PerkinElmer). PF74 and lenacapavir (LEN) are HIV-1 capsid inhibitors. N/A, not available.

**Table 2.** Anti-HIV-1 activity of ACAi-001 against HIV-1, HIV-2, or HIV-1 capsid inhibitor (BVM)-resistant variant.

| Virus | Cells | EC<sub>50</sub> ($\mu\text{M}$) | | | |
| --- | --- | --- | --- | --- | --- |
|  |  | ACAi-001 | ABC | 3TC | PF74 |
| HIV-1<sub>LAI</sub> | MT-2 | 0.5 ± 0.1 | 3.3 ± 0.1 | 2.9 ± 0.3 | 0.34 ± 0.05 |
| HIV-2<sub>ROD</sub> | MT-2 | 1.6 ± 0.3 | 4.0 ± 0.9 | >10 | >10 |
| HIV-1<sub>BVM</sub><sup>R</sup> | MT-4 | 2.0 ± 0.3 | >10 | 4.2 ± 0.1 | >10 |

MT-2 cells ( $10^4/\text{mL}$ ) were exposed to $100\times$ TCID<sub>50</sub> of HIV-1<sub>LAI</sub> or HIV-2<sub>ROD</sub> in the presence of various compound concentrations, and EC<sub>50</sub> values were determined by MTT assay. For HIV-1<sub>BVM</sub><sup>R</sup> (a bevirimat-resistant variant generated by drug selection), MT-4 cells ( $10^5/\text{mL}$ ) were exposed to $100\times$ TCID<sub>50</sub> and EC<sub>50</sub> values were determined based on inhibition of p24 Gag production. All assays were performed in duplicate; data represent mean ± SD from two or three independent experiments. Control compounds: abacavir (ABC) and lamivudine (3TC; RT inhibitors), and PF74 (capsid inhibitor).

**Table 3.**
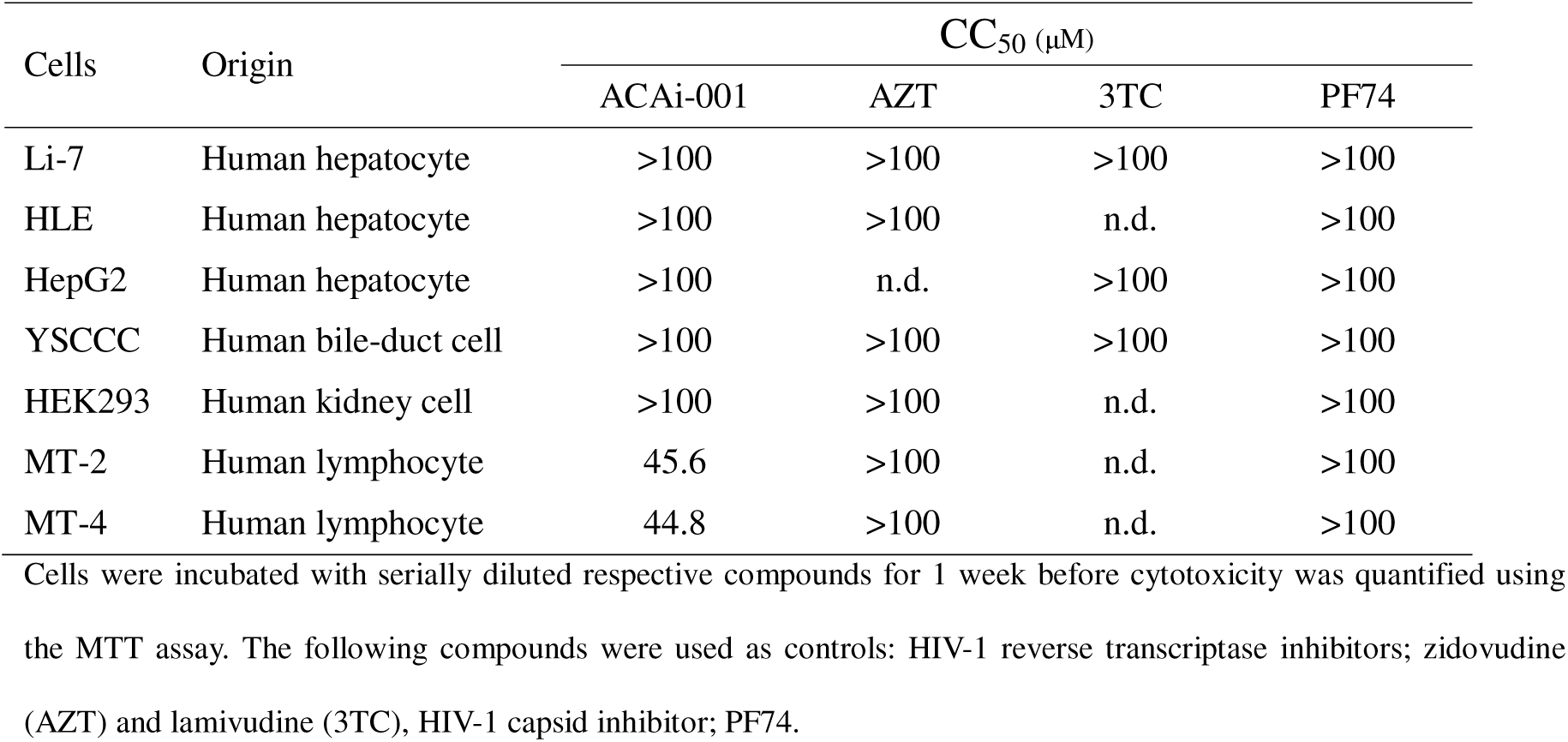
Cytotoxicity of ACAi-001 against various human cell-lines.

### ACAi-001 binding covalently to CA induces CA multimerization and degradation

To investigate the direct binding of ACAi-001 to CA, we purified CA proteins expressed in *E. coli* (Figure S2A). Evaluation using liquid chromatography-mass spectrometry (LC-MS) showed that one, two, and three ACAi-001 molecules bound to the CA monomer under native conditions (Figure S2B). One and two ACAi-001 molecules were found to interact with the CA monomer under denaturing conditions involving treatment with acetonitrile and trifluoroacetic acid (Figure 2A), suggesting that ACAi-001 directly and covalently binds to CA proteins similar to ebselen (Eb), a covalent capsid inhibitor (14). The MW of the ACAi-001 plus CA monomer complex detected by LC-MS was five units smaller than the calculated value, suggesting covalent bond formation accompanied by the loss of small molecules (Figure S2C). In addition, CA proteins were incubated with ACAi-001, ACAi-028, the HIV-1 integrase (IN) inhibitor raltegravir (RAL) (15), and the capsid inhibitor PF74 (16), which is a lead compound of lenacapavir (17), for 72 h in reaction buffer. The results showed that large amounts of CA multimers and degradation products induced by ACAi-001 were detected by western blotting (WB) with an anti-CA antibody (Figure 2B) under denaturing conditions compared to the effects of ACAi-028, RAL, and PF74. CA multimerization began to occur after 3 h in the presence of 100 μM of ACAi-001, whereas CA degradation products appeared after 12 h (Figure 2C). These aberrant CA products were detected at ACAi-001 concentrations exceeding 20 μM (Figure 2D). When the samples were treated at different temperatures, CA multimerization and degradation were induced at higher temperatures (Figure S2D). Moreover, CA proteins spanning a wide range of molecular sizes, including the CA monomer, appeared as broad, heterogeneous peaks 24 h after ACAi-001 treatment using size-exclusion chromatography (SEC) (Figures 2E and S2E). To determine whether the covalent interactions of ACAi-001 induce such aberrant multimerization and degradation of other proteins, we confirmed the effect of ACAi-001 on human serum albumin (HSA) and bovine serum albumin (BSA). When HSA and BSA were analyzed using WB after treatment with dimethyl sulfoxide (DMSO) or ACAi-001 for 72 h, no abnormal HSA and BSA multimerization or degradation was observed in ACAi-001-treated samples, compared to that observed for CA proteins (Figure S2F). Next, we examined the profiles of CA multimerization (18) and thermal stability (19) after treatment with ACAi-001 (Figures 2F and 2G). Interestingly, ACAi-001 (>20 μM) impaired normal, sodium-dependent CA multimerization after 24 h, unlike RAL, indicating that ACAi-001-induced aberrant multimerization renders CA incapable of proper self-assembly (Figure 2F). Consistent with this loss of physiological function, ACAi-001 also markedly decreased CA thermal stability, by 8.4°C and 12.4°C at 20 and 200 μM, respectively, after 72 h of treatment (Figure 2G).

**Figure 2.**
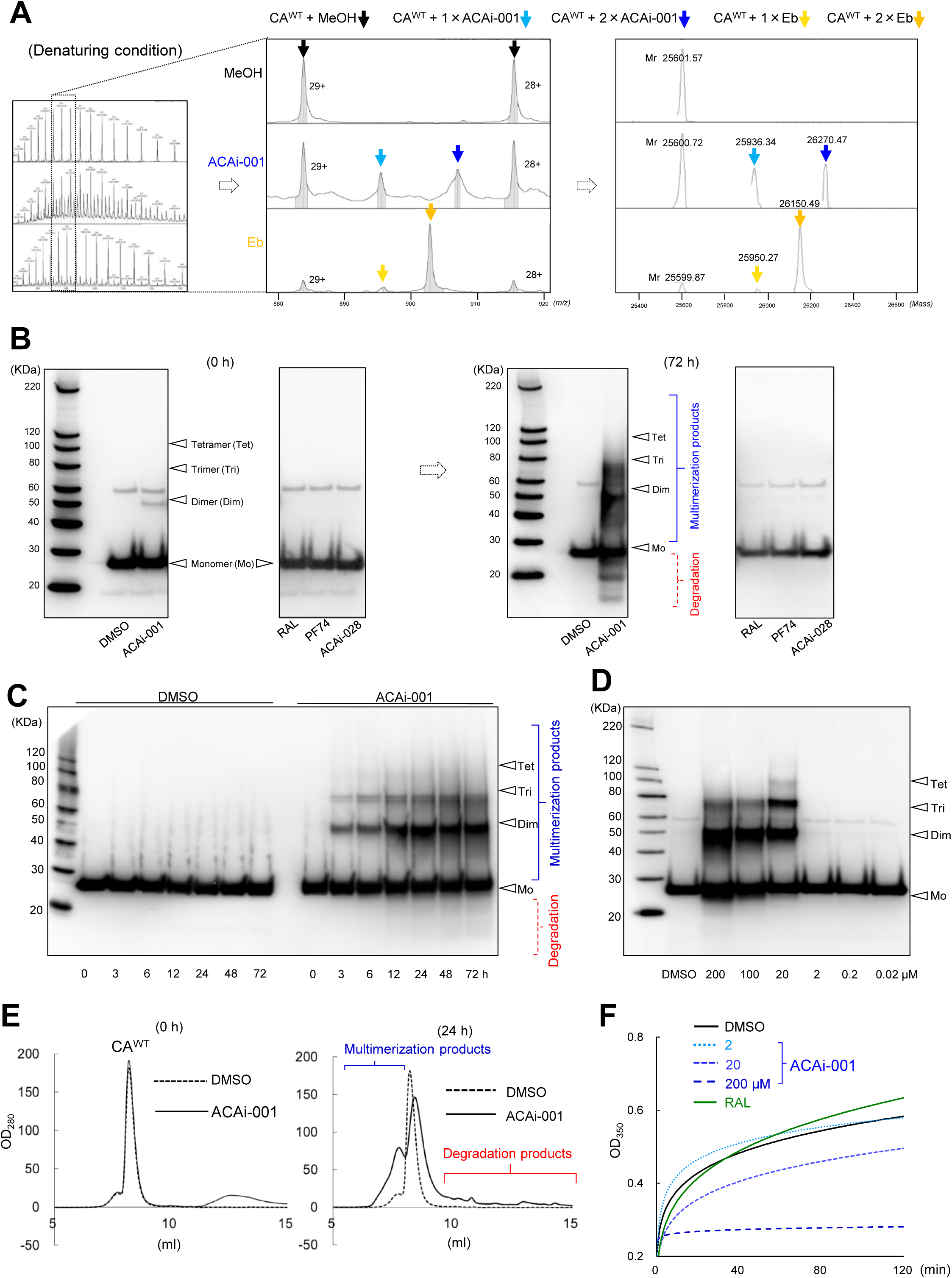
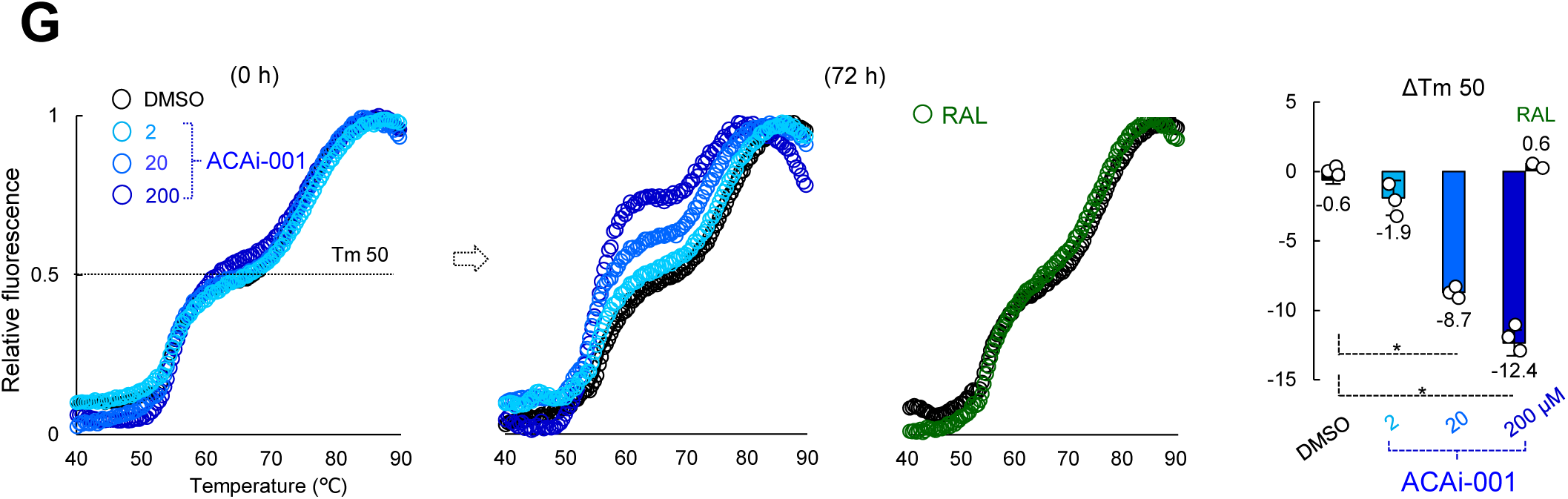
CA multimerization and degradation profile induced by ACAi-001. (**A**) Native LC-MS deconvolution (right) of CA monomer (10 µM) charge-state peaks (black arrows) with one or two bound ACAi-001 (50 µM; light and dark blue arrows) or Eb (50 µM; yellow and orange arrows), analyzed under denaturing conditions. (**B**) WB of CA treated with DMSO (1%), ACAi-001 (100 µM), RAL (100 µM), PF74 (20 µM), or ACAi-028 (20 µM) for 0 h (left) or 72 h (right); arrows indicate monomer (Mo), dimer (Dim), trimer (Tri), and tetramer (Tet). (**C**, **D**) Time- (0–72 h) and dose- (0.02–200 µM) dependent aberrant CA multimerization and degradation induced by ACAi-001, assessed by WB relative to DMSO. (**E**) SEC profiles of CA treated with DMSO or ACAi-001 (100 µM) for 0 h (left) or 24 h (right). (**F**) CA multimerization assay: high-sodium buffer was added to CA pre-incubated for 24 h with ACAi-001 (2, 20, or 200 µM) or RAL (100 µM) under low-sodium conditions; turbidity (OD_350_) was monitored for 120 min. (**G**) Differential scanning fluorimetry using SYPRO Orange with DMSO, ACAi-001 (2, 20, or 200 µM), or RAL (100 µM); ΔTm over 0–72 h is shown (right). All assays were performed in triplicate (≥3 independent experiments); error bars, ±SD. *, *P* < 0.05 (Student’s t-test).

### CA multimerization and degradation induced by ACAi-001 are readily detected by WB and p24 enzyme-linked immunosorbent assay

To detect CA multimerization and degradation induced by the selected compounds, the states of CA were evaluated and measured as the CA immunogenicity using WB and an automated enzyme-linked immunosorbent assay (ELISA) device (Lumipulse G1200). We constructed plasmids expressing wild-type HIV-1_NL4-3_ CA (CA^WT^). Cell lysates containing CA^WT^ were incubated with ACAi-001 for different times (0, 24, 48, and 72 h) at 37°C and then evaluated for CA immunogenicity. Interestingly, the immunogenicity of CA monomers treated with ACAi-001 decreased significantly in a time-dependent manner and was less than 20% within 72 h of incubation compared to that in the DMSO control (Figures 3A and S3A), suggesting that ACAi-001 induces aberrant multimerization and degradation of CA derived from the cell lysates. In contrast, the amount of CA^WT^ was not significantly reduced when cell lysates containing CA^WT^ incubated with DMSO and PF74 were used as negative controls (Figure 3B). Next, we examined the profiles of CA treated with ACAi-001 using lysates of infectious HIV-1_NL4-3_ (HIV^WT^) by ELISA and WB analyses. Figures 3C and 3D show that ACAi-001 also significantly reduced CA immunogenicity in the lysate containing HIV^WT^ viral lysates. CA immunogenicity as measured by ELISA was reduced to less than 20% within 24 h of ACAi-001 treatment, whereas only a minor reduction was observed with DMSO. We evaluated the effects of the first-in-class HIV-1 CA maturation inhibitors bevirimat (20) and PF74 on CA. As shown in Figure 3D, bevirimat and PF74 did not reduce CA immunogenicity, suggesting that ACAi-001 exerts a different mechanism(s) of interaction with CA than that of other HIV-1 CA inhibitors. Figure 3E shows that the CA immunogenicity induced by ACAi-001 was reduced in a dose-dependent manner. Additionally, ACAi-001 reduced HIV-2_ROD_ (37) CA immunogenicity, even in the lysates of HIV-2_ROD_-infected MT-2 cells (Figure 3F). Furthermore, ACAi-001 induced CA degradation as assessed using cell lysates from peripheral blood mononuclear cells infected with four different subtypes of clinically isolated HIV-1 strains (Figure S3B). These results indicate that ACAi-001 can induce multimerization and degradation of recombinant CA protein expressed in the cell lysates and CA in HIV^WT^ particles, which were readily detected using WB and p24 ELISA.

**Figure 3.**
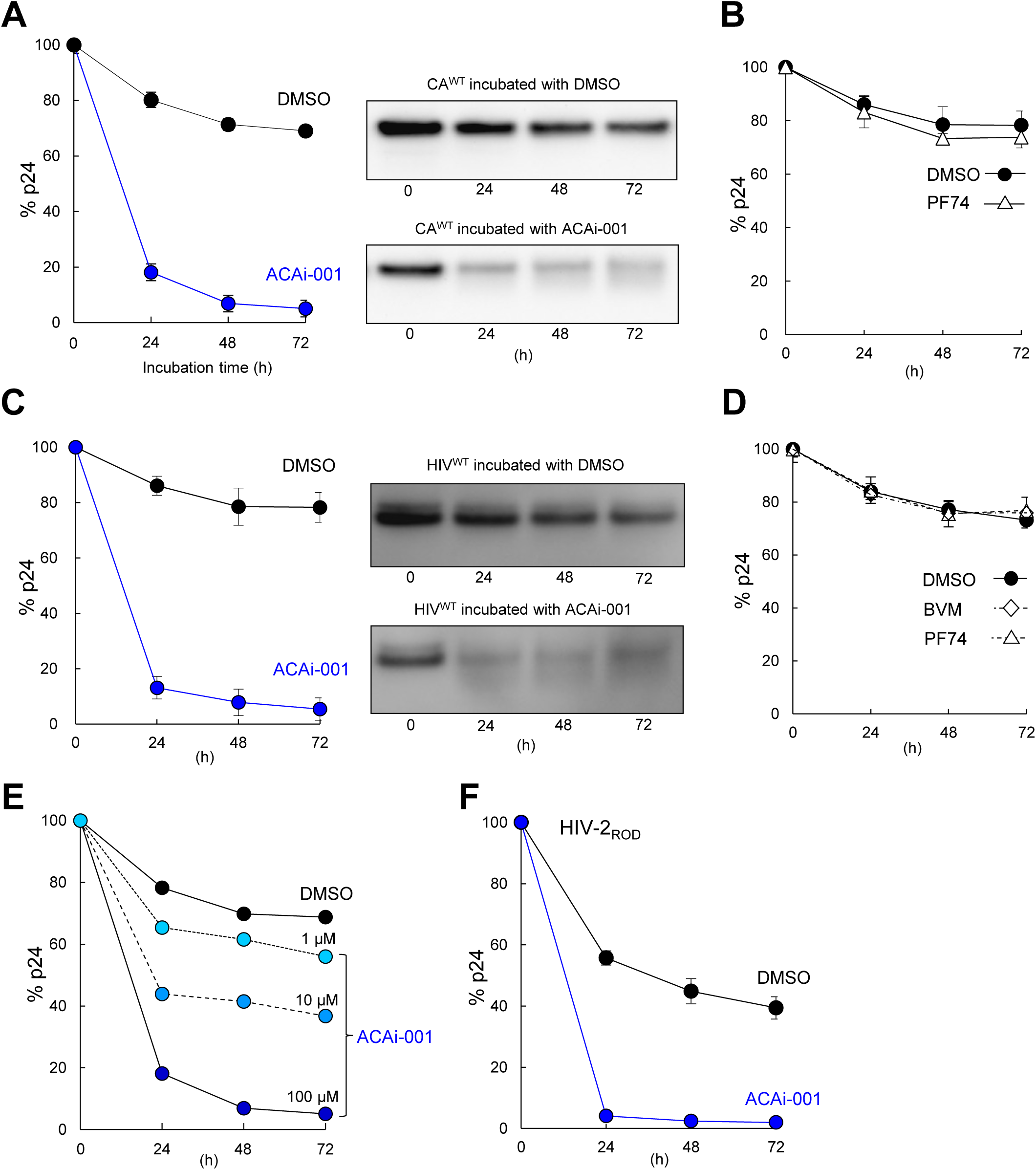
Detection of ACAi-001-induced CA multimerization and degradation in cellular expression lysates. (**A**) CA cell lysates treated with DMSO or ACAi-001 (blue) for 24–72 h at 37°C, assessed by automated p24 ELISA (left) and WB (right). (**B**) Lysates treated with DMSO or PF74, assessed by ELISA. (**C**) HIV-1_NL4-3_ lysates treated as in (**A**), assessed by ELISA (left) and WB (right). (**D**) Lysates treated with DMSO, bevirimat (BVM), or PF74. (**E**) Dose-dependent effects of ACAi-001 (1, 10, or 100 µM) on CA cell lysates at 37°C, by ELISA. (**F**) HIV-2_ROD_ lysates treated with DMSO or ACAi-001 for 24–72 h at 37°C, by ELISA.

### Putative covalent bond-forming structures in ACAi-001

ACAi-001 has a hydroquinone moiety that is capable of forming covalent bonds with CA proteins (Figure 4A) (21). The MW of the ACAi-001 CA monomer complex detected by LC-MS was approximately 5.0 units lower than the calculated value (Figure S2C). We hypothesized that ACAi-001 forms covalent linkages with CA proteins through a mechanism requiring at least two sequential oxidation events (Figure 4A). In the first oxidation step, the hydroquinone moiety of ACAi-001 is converted to a p-benzoquinone, which likely forms a covalent bond with a nucleophilic amino acid residue such as cysteine and serine. The resulting adduct can then be re-oxidized to regenerate the p-benzoquinone form, enabling a second covalent bond formation with another CA protein. Through these two oxidation steps, a single ACAi-001 molecule is capable of crosslinking two CA proteins, thereby acting as a molecular hinge (Figure 4A), which may ultimately induce aberrant CA multimerization. Through this mechanism, the theoretical MW of the ACAi-001 and CA monomer complex decreases by 4.0 units, leading to close agreement with the value observed by LC-MS. To examine these reactions between CA proteins and ACAi-001, we added the reducing agent β-mercaptoethanol (βME) to the reaction buffer to prevent oxidation of the hydroquinone. As shown in Figures 4B and S4, aberrant CA multimerization was found in the normal reaction buffer after treatment with more than 20 μM of ACAi-001 for 24 h, whereas such aberrant CA multimerization was completely inhibited by the addition of 10 mM βME. In addition, ANS-1028 and ANS-1050 are structural analogs of ACAi-001 in which the hydroquinone moiety is replaced by a meta- or ortho-hydroxyphenyl group, eliminating one hydroxyl group from the benzene ring (Figure 4C). As expected, ANS-1028 and ANS-1050 did not cause CA multimerization and degradation, nor did PF74 and Eb (Figure 4C and Table S2). These results indicate that the hydroquinone structure of ACAi-001 is critical for the formation of covalent bonds in CA, which induce aberrant CA multimerization and degradation.

**Figure 4.**
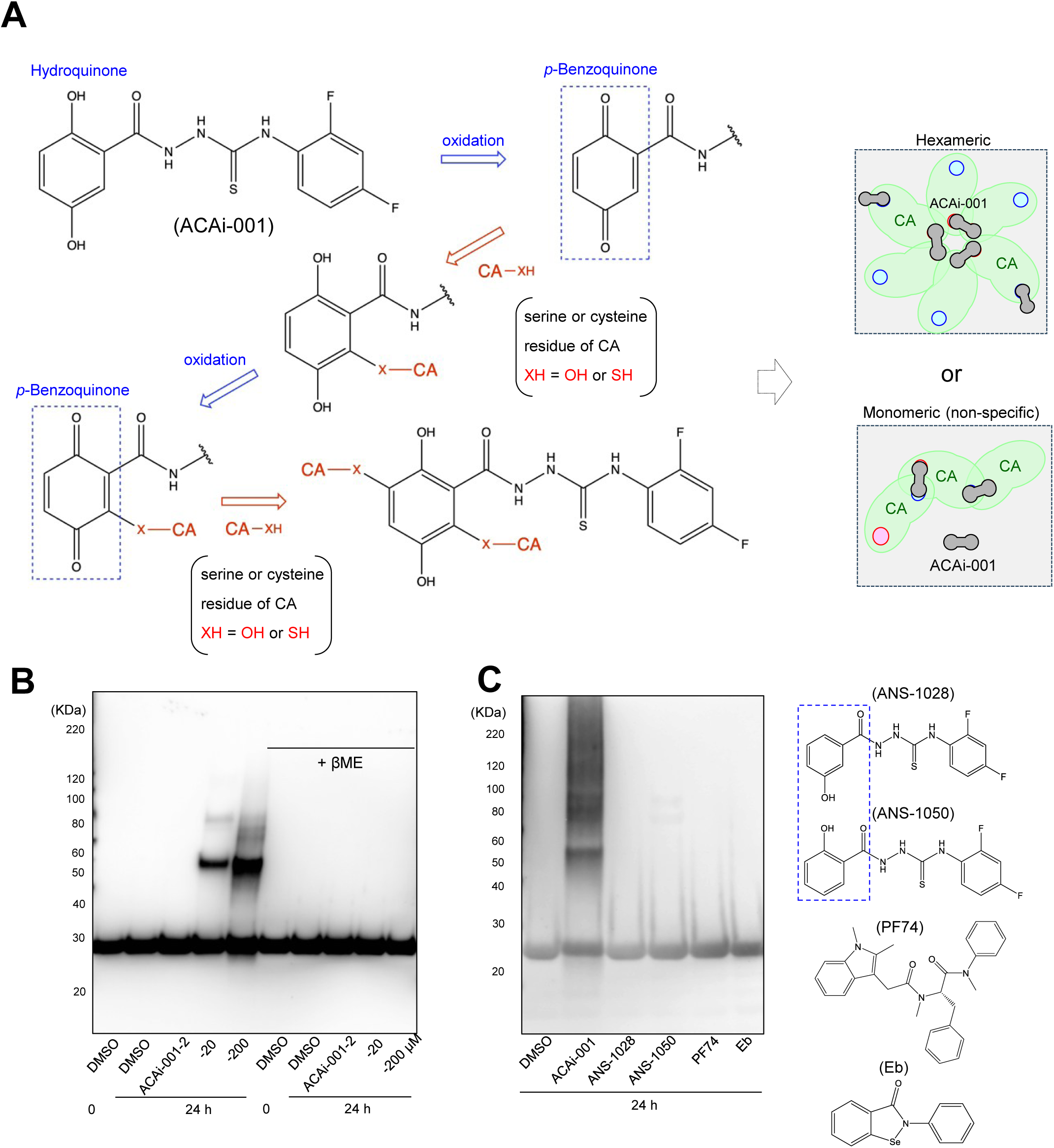
Putative covalent reactions of ACAi-001 to CA proteins. (**A**) Proposed mechanism of ACAi-001 reactivity toward CA. Oxidation converts the hydroquinone moiety of ACAi-001 to p-benzoquinone (dashed blue square), which forms a covalent bond with a nucleophilic residue (e.g., cysteine or serine) in CA. Re-oxidation of the resulting adduct regenerates the p-benzoquinone form, enabling a second covalent bond with an adjacent CA protein; through these two sequential oxidation steps, a single ACAi-001 molecule crosslinks two CA proteins. The gray box schematizes the resulting aberrant CA multimerization in hexameric and monomeric states. (**B**) Dose-dependent aberrant CA multimerization and degradation induced by ACAi-001 (2, 20, and 200 µM) after 24-h incubation in reaction buffer without (left) or with (right) the reducing agent βME (10 mM). (**C**) WB (anti-p24 antibody) of CA treated with ACAi-001, ANS-1028, ANS-1050, PF74, or Eb (100 µM each). Chemical structures are shown on the right; the dashed blue square marks the phenol formed when the hydroquinone of ACAi-001 is replaced by a single hydroxyl group in ANS-1028 and ANS-1050. All assays were performed independently at least twice; representative data are shown.

### ACAi-001 binds to a serine residue in the target cavity of CA-NTD and cysteine residues in CA-CTD

To investigate how ACAi-001 interacts with specific AAs and induces CA multimerization and degradation, we constructed a plasmid expressing CA truncated at the N-terminal AAs (CA-NTD^Δ1-100^). As shown in Figure S5A, the cell lysates of COS7 cells expressing CA-NTD^Δ1-100^ incubated for 72 h with ACAi-001 showed no significant structural changes, similar to the DMSO-treated control, suggesting that ACAi-001 requires full-length CA to induce CA multimerization and degradation.

ACAi-001, carrying an electrophilic reactive group, hydroquinone, can covalently and irreversibly bind to nucleophilic AA residues in the target CA protein. We predicted that ACAi-001 has the potential to form covalent bonds with some cysteine and serine residues of CA (9, 21). To distinguish the putative ACAi-001 binding cavities, we designated the target cavity in the NTD as the "ACAi-pocket" and the harboring two cysteines in the C-terminal domain as the "Cys-pocket", as shown in Figure 5A. To confirm whether ACAi-001 interacts with the AA residues in these target cavities (Figure 5A), we produced CA variants carrying S16E (CA^S16E^) in the ACAi-pocket, C198A/C218A (CA^C198A/C218A^) in the Cys-pocket, or S16E/C198A/C218A (CA^S16E/C198A/C218A^) in both pockets (Figure S5B), which may alter the binding ability of ACAi-001 to CA. Sequence analysis of 12,261 HIV-1 sequences (all subtypes; Los Alamos HIV Sequence Database) using WebLogo 3 confirmed high conservation of Ser16, Cys198, and Cys218 (Figure 5A). SEC analysis showed aberrant CA^WT^ multimerization with two broad peaks and CA^WT^ degradation on a gentle slope after treatment with ACAi-001 for 24 h, compared with those in the DMSO control (Figure 5B). Although the peaks of CA^S16E^ and CA^C198A/C218A^ multimerization and degradation induced by ACAi-001 treatment decreased, the peaks of CA^S16E/C198A/C218A^ induced by ACAi-001 were similar to those following DMSO treatment (Figure 5B). In line with the SEC results, CA^WT^ multimerization was induced in a dose-dependent manner by ACAi-001 (5, 10, and 20 μM) under denaturing conditions according to WB, whereas CA^S16E^ multimerization and CA^C198A/C218A^ multimerization were slightly and strongly decreased after treatment with ACAi-001 for 24 h. The multimerization of CA^S16E/C198A/C218A^ carrying these three mutations induced by ACAi-001 almost disappeared, as shown in Figure 5C. In addition, binding of ACAi-001 to CA^S16E/C198A/C218A^ significantly decreased under LC-MS denaturing conditions. The binding peaks of the calculated MW of CA^S16E/C198A/C218A^ plus ACAi-001 disappeared, except for a negligible peak of one ACAi-001 binding event (Figures 5D, S5C, and S5D). These results suggest that ACAi-001 covalently interacts with Ser16 in the ACAi-pocket and Cys198/Cys218 in the Cys-pocket *in vitro*.

**Figure 5.**
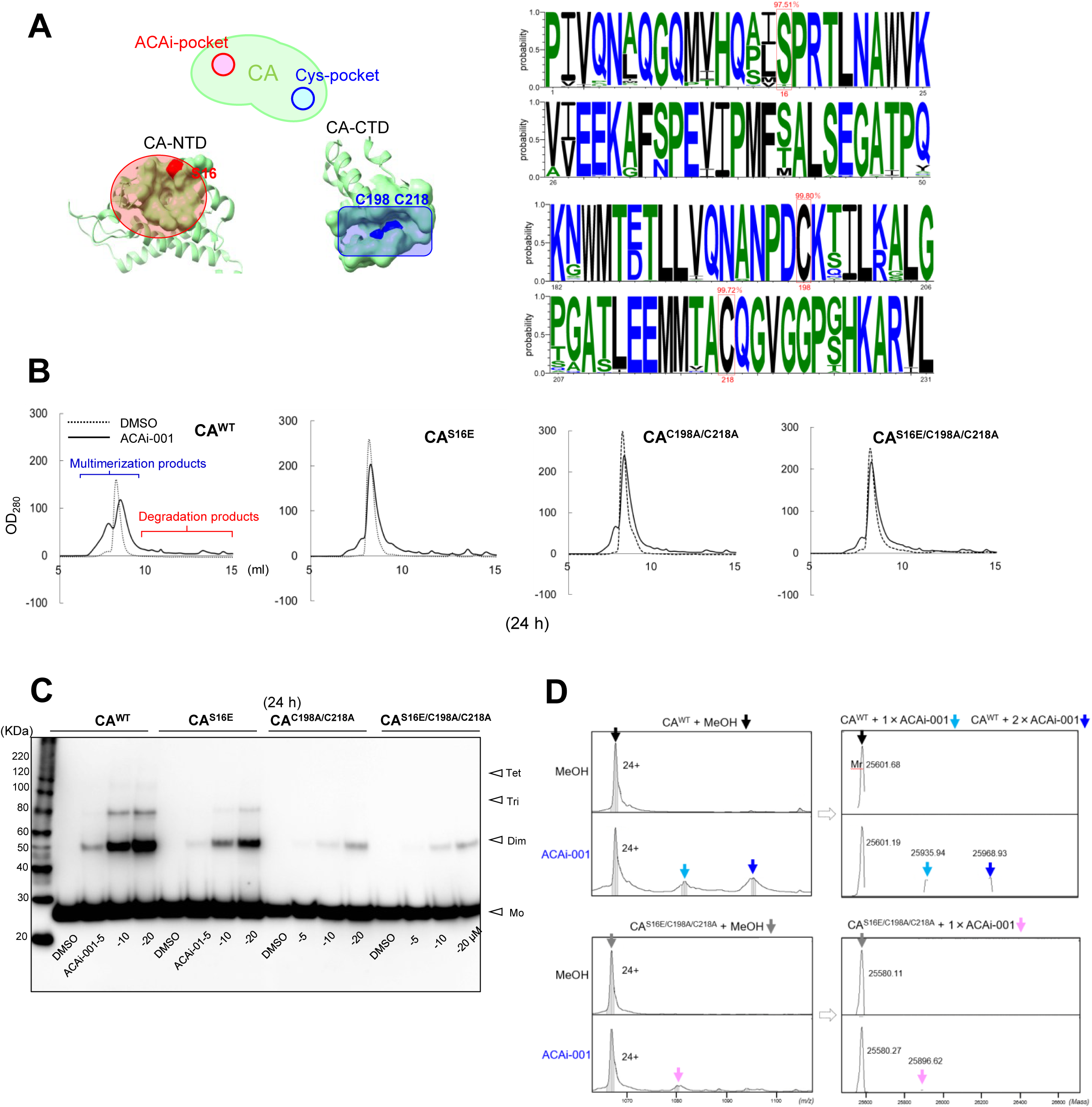
ACAi-001 interacts with Ser16 in the CA-N-terminal domain (NTD) and Cys198 and Cys218 in the CA-C-terminal domain (CTD). (**A**) Ser16 (red) lies within the ACAi-pocket of the CA-NTD; Cys198 and Cys218 (blue) lie within the Cys-pocket of the CA-CTD. WebLogo 3 visualization of amino acid frequencies in the CA-NTD (residues 1– 50) and CA-CTD (residues 182–231), generated from 12,261 HIV-1 sequences across all subtypes (Los Alamos HIV Sequence Database, 2022 Web alignment); Ser16, Cys198, and Cys218 (red boxes) show percentage consensus (Jalview 2.11.1.4). (**B**) SEC analysis of CA^WT^, CA^S16E^, CA^C198A/C218A^, and CA^S16E/C198A/C218A^ multimers induced by ACAi-001 (100 μM) after 24-h incubation. (**C**) WB of aberrant CA multimerization in CA^S16E^, CA^C198A/C218A^, and CA^S16E/C198A/C218A^ variants treated with ACAi-001 (5, 10, or 20 μM); arrows indicate theoretical MW of monomer (Mo), dimer (Dim), trimer (Tri), and tetramer (Tet). (**D**) LC-MS (denatured state) of ACAi-001 binding to CA^WT^ and CA^S16E/C198A/C218A^; deconvoluted MWs are shown (right). All assays were performed independently at least twice; representative data shown.

### Effect of ACAi-001 on the early- and late-stage of the HIV-1 life cycle

We investigated the time point at which the anti-HIV-1 activity of ACAi-001, which induces CA multimerization and degradation, is exerted during the HIV-1 life cycle. To examine early-stage inhibition by ACAi-001 (Figure 6A), we performed TZM-bl cell and colorimetric reverse transcriptase (RT) activity assays. When ACAi-001 was added simultaneously with VSV-G HIV-1_dENV_ to TZM-bl cells, it did not significantly inhibit early-stage HIV-1 infection (Figure 6B). Similar results were obtained with the protease inhibitor DRV (22), in contrast to the RT inhibitor AZT (23) and RAL, both of which potently suppressed infection under these conditions (Figure 6B). ACAi-001, DMSO, and PF74 did not significantly impact HIV-1 RT activity compared to the effects of a non-nucleoside RT inhibitor, efavirenz (24) (Figure 6C). To confirm the specificity of ACAi-001, we examined its effects on HIV-1 RT and IN using viral particles, and on HTLV-1 Gag proteins using HTLV-1-infected MT-2 cells. After incubation with ACAi-001 (100 µM) for 72 h at 37°C, WB analysis showed RT, IN, and HTLV-1 Gag (p19, p24, p53) levels comparable to the DMSO control, indicating that ACAi-001 does not affect these off-target proteins (Figures S6A and S6B). Together, these results indicate that ACAi-001 selectively targets CA without affecting other retroviral proteins, supporting its specificity as a CA-targeted covalent inhibitor.

**Figure 6.**
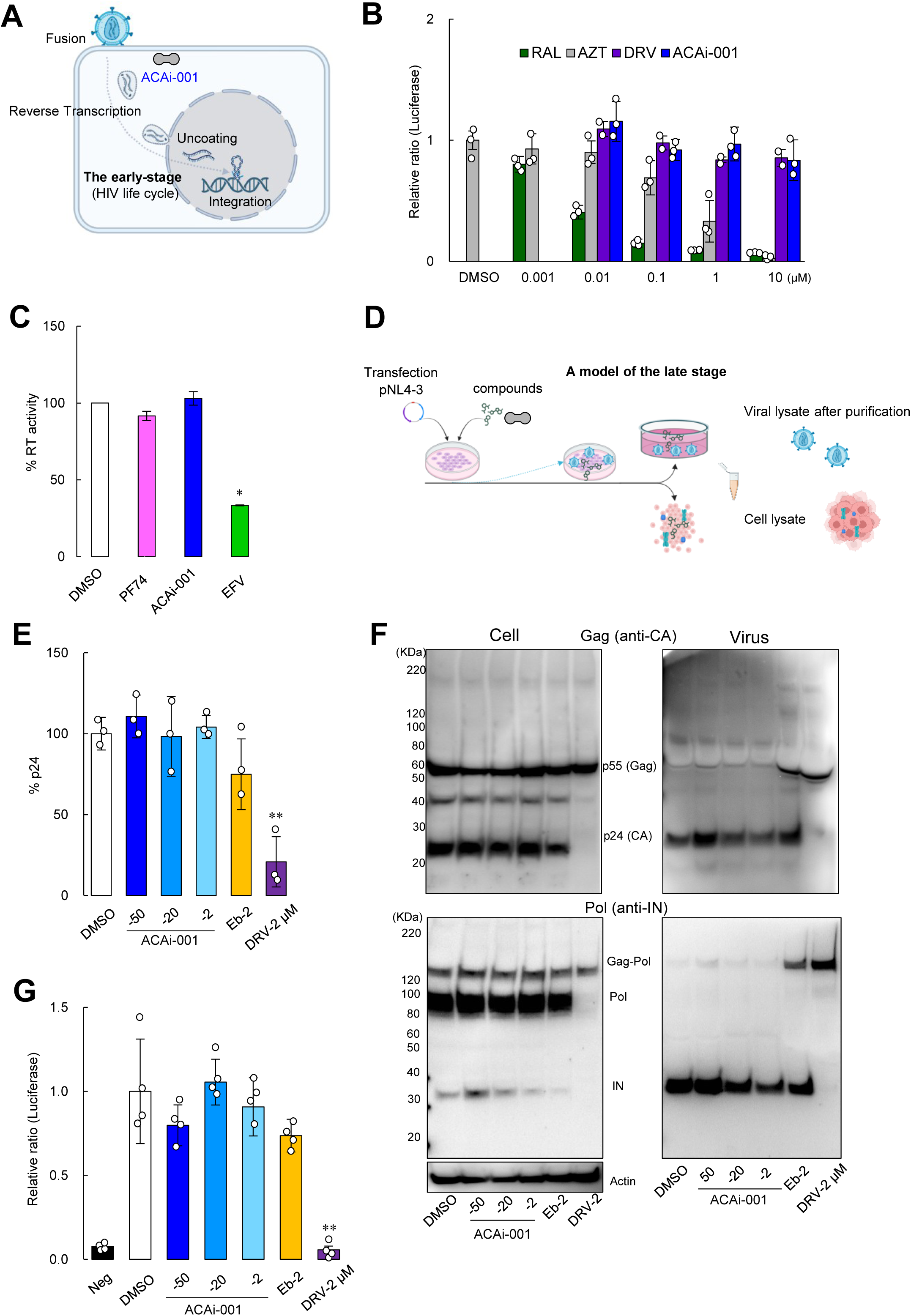
Anti-HIV-1 effects of ACAi-001 on the HIV-1 life cycle. (**A**) Schematic of the early-stage HIV-1 life cycle. (**B**) Early-stage inhibition of ACAi-001 (blue), RAL (dark green), AZT (grey), and DRV (purple) against VSV-G HIV-1_ΔENV_, determined by TZM-bl assay. (**C**) RT activity in the presence of DMSO (0.5%), ACAi-001 (100 μM), PF74 (100 μM), or EFV (2 μM), determined by colorimetric RT assay; OD_405_ is expressed as percent RT activity relative to DMSO (mean ± SD, 2 independent experiments in duplicate). (**D**) Schematic of the late-stage HIV-1 life cycle. (**E**) Virus production, determined by p24 levels in the supernatant of pNL4-3-transfected 293T cells treated with ACAi-001 (50, 20, or 2 µM), Eb (2 µM), or DRV (2 µM). (**F**) WB of Gag and Gag-Pol (anti-Gag and anti-IN antibodies) and actin (loading control) in transfected-cell lysates, and of Gag-Pol processing and maturation in partially purified virions from the culture supernatant. (**G**) Infectivity of the partially purified virions, determined by TZM-bl assay. All experiments were performed independently 2–3 times; representative data shown. Error bars, ±SD.

Next, to investigate the effect of ACAi-001 on late-stage inhibition (Figure 6D), we determined whether ACAi-001 affected HIV-1 production, including Gag and Gag-pol proteolytic processing and maturation. At 48 h after transfection of pHIV-1_NL4-3_ into 293T cells in the presence of ACAi-001 (50, 20, and 2 µM), Eb (2 µM), or DRV (2 µM), viral production was assessed using ELISA and WB to compare the p24 expression levels in the culture supernatant and those of the intracellular Gag and Gag-pol proteins (Figure 6E). ACAi-001 and Eb did not significantly affect viral production, whereas DRV treatment reduced HIV-1 production (Figure 6E). Unlike DRV, ACAi-001 and Eb did not affect Gag proteins in the cells, as shown on the left side of Figure 6F. Additionally, the effect of ACAi-001 on Gag proteolytic processing and maturation of HIV-1 virions was investigated using WB and a TZM-bl assay after virus purification. ACAi-001 did not affect HIV-1 maturation, as observed via WB with anti-Gag and anti-IN antibodies, similar to the results observed in the DMSO control, as shown in the right panel of Figure 6F. Furthermore, viruses produced in the presence of DMSO, ACAi-001, or Eb did not significantly reduce the infectivity of TZM-bl cells, in contrast to the effects of DRV (Figure 6G). Eb binds to cysteines in the CA-CTD and has been reported to inhibit the early-stage, but not the late-stage, of HIV-1 replication (13, 14). Interestingly, incompletely processed (immature) products such as Gag-pol (p160), Gag (p55), and Gag intermediates were found in purified viruses produced at 2 µM of Eb according to WB with anti-Gag or anti-IN antibodies (Figure 6F), whereas the infectivity of virions containing incompletely processed Eb products was not significantly reduced (Figure 6G). These results suggest that Eb affects Gag-pol processing without inhibiting the late-stage of the HIV-1 life cycle. Additionally, ACAi-001 did not significantly impact the early and late stages of the HIV-1 life cycle, in contrast to capsid inhibitors such as Eb and PF74 (16).

#### ACAi-001 inhibits HIV-1 infection through direct action on viral particles

CA proteins constitute the capsid lattice (HIV-1 core) comprising approximately 250 hexamers and 12 pentamers (11), with the target cavity located inside the CA hexamer (Figure 7A). To investigate the location and dimensions of the binding pocket within the CA hexamer, we constructed CA hexamer models in both the presence and absence of ACAi-001, based on the CA crystal structure (PDB ID: 8CKV). Our model indicates that ACAi-001 binds to a pocket within the CA-NTD interface formed by a CA dimer (Figure 7A). Furthermore, docking simulations predicted the binding profile of ACAi-001 in the hexameric state: the compound putatively forms two hydrogen bonds (H-bonds) with Q13 and S16 of one CA monomer, and a third H-bond with S16 of the adjacent monomer (Figure 7A), suggesting that it initially bridges adjacent CA monomers before forming covalent bonds within the hexameric HIV-1 core. Treatment of CA hexamers with ACAi-001 promoted greater CA multimerization at high (3 M) than at low (150 mM) NaCl concentration (Figure 7B), and in the presence of IP-6, which stabilizes CA hexamers (38), higher-order CA multimers formed at a lower ACAi-001 concentration (25 µM) than without IP-6 (Figure S7A), indicating that ACAi-001 preferentially promotes CA multimerization under conditions favoring hexameric assembly.

**Figure 7.**
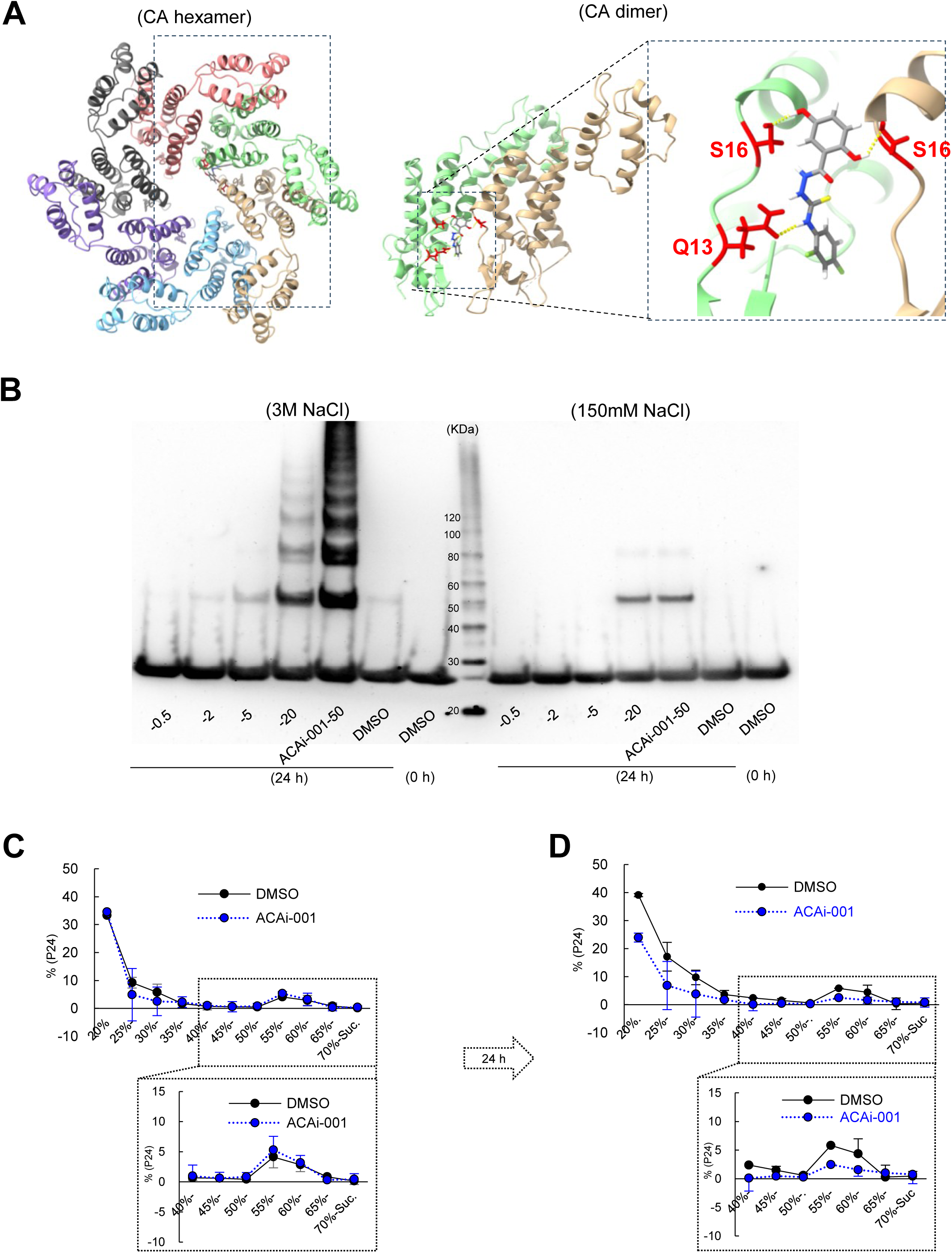
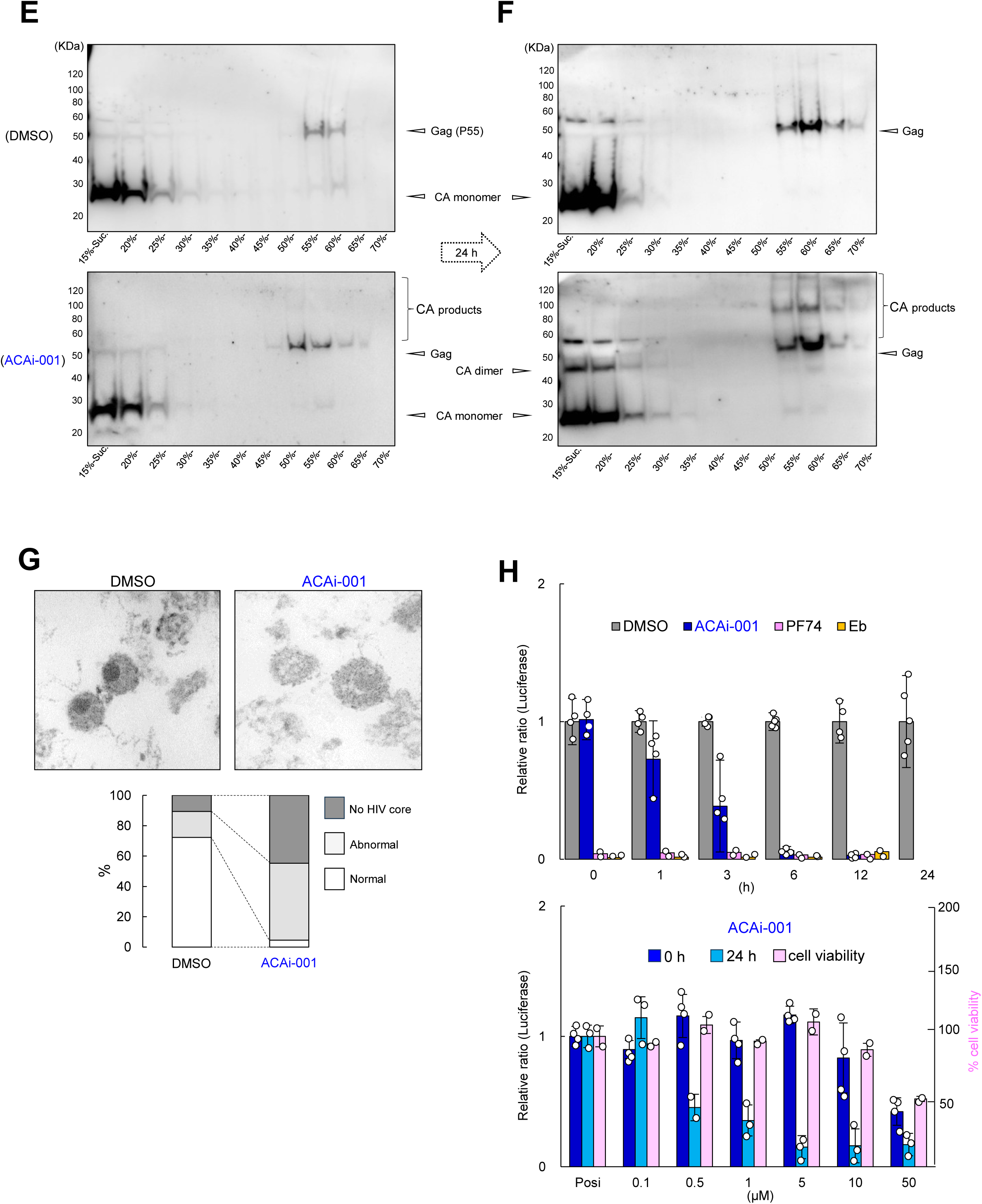
ACAi-001 directly and covalently targets HIV-1 viral particles (CA hexamer). (**A**) Docking model of ACAi-001 bound to a CA dimer within the hexameric lattice, forming putative hydrogen bonds with Q13 and S16 of one monomer and S16 of the neighboring monomer. (**B**) WB of CA multimerization after 24-h incubation with ACAi-001 (0.5–50 µM) or DMSO in high-(3 M) or low-salt (150 mM) sodium buffer. (**C-F**) Sucrose density gradient fractionation (20–70%) of purified HIV-1 viral cores after treatment with DMSO or ACAi-001 for 0 h (**C, E**) or 24 h (**D, F**), analyzed by ELISA (**C, D**) and WB (**E, F**) with anti-CA antibody. (**G**) TEM of purified viruses treated with DMSO or ACAi-001 (100 µM) for 72 h at 37°C, with viral morphology (normal, abnormal, or no core) compared as a ratio. (**H**) TZM-bl infectivity of VSV-G HIV-1_ΔENV_ pre-incubated with DMSO, ACAi-001 (50 µM), PF74 (10 µM), or Eb (10 µM) for 0–24 h, and dose-dependent inhibition by ACAi-001 (0.1–50 µM) following 0 h (dark blue) or 24 h (light blue) pre-incubation; TZM-bl cell viability at each ACAi-001 concentration is also shown. Assays were performed 2–3 times independently; representative data shown; error bars, ±SD.

Purified HIV-1 particles were treated with DMSO or ACAi-001 for 24 h. The envelopes of these viruses were removed using Triton X-100, and the envelope-stripped cores were enriched by ultracentrifugation through a continuous 20–70% sucrose gradient, as previously reported (25, 26). After ACAi-001 treatment for 24 h, the CA immunogenicity peak, including the HIV-1 core from 50–65% sucrose gradient fractions by ELISA, was decreased compared to that following DMSO treatment (Figures 7C and 7D). We also confirmed the state of CA in these sucrose gradient fractions by denaturing WB using a CA anti-polyclonal antibody. In the presence of DMSO, CA monomers were detected in the low-density sucrose fractions, while p55 (Gag) was found in the higher-density HIV-1 core fractions (upper panel of Figure 7C). Following 24 h of DMSO treatment, CA monomers and Gag were detected in the same levels as observed prior to treatment (upper panel of Figure 7D). In contrast, upon addition of ACAi-001, CA and Gag products were faintly detected in the HIV-1 core fractions compared to after DMSO treatment (lower panel of Figure 7E). Furthermore, following 24 h of ACAi-001 treatment, Gag and various CA products, including aberrant multimerization, were clearly observed in the HIV-1 core fractions (lower panel of Figure 7F). Consistently, direct time-dependent treatment of viral particles with ACAi-001 also induced multimerization of CA (Figure S7B). These results indicate that ACAi-001 can directly and covalently bind CA and affect the HIV-1 core in viral particles. Interestingly, when the morphology of mature HIV-1_NL4-3_ virions was observed using transmission electron microscopy (TEM) after incubation with ACAi-001 for 72 h, the HIV-1 core in the viral particle was remarkably decomposed by ACAi-001 (4.5% normal, 49.2% abnormal, and 44.8% no HIV-1 core structures) compared with the effects of DMSO treatment (72.3% normal, 17.1% abnormal, and 10.6% no HIV-1 core structures) (Figure 7G) (26), indicating that ACAi-001 penetrates the viral envelope and directly disrupts the HIV-1 core.

Finally, we determined the time required for ACAi-001 to exert anti-HIV-1 activity against viral particles by pre-incubating HIV-1 particles with ACAi-001, PF74, or Eb for 0, 1, 3, 6, 12, or 24 h before infecting TZM-bl cells. PF74 and Eb inhibited infectivity without pre-incubation, whereas ACAi-001 required at least 3 h (Figures 7H and S7D), consistent with the onset of aberrant CA multimerization observed at 3 h by WB (Figure 2C).

Collectively, these results indicate that ACAi-001 directly and covalently binds CA within the HIV-1 core of viral particles, inducing time-dependent aberrant CA multimerization and degradation that decomposes the core and is associated with the direct anti-HIV-1 activity of ACAi-001 against HIV-1 particles.

## Discussion

We identified ACAi-001 as a covalent anti-HIV-1 CA inhibitor that binds the ACAi-pocket in the CA-NTD, in addition to the Cys-pocket in the CA-CTD (Figure 5A). We propose that ACAi-001 engages the ACAi-pocket within the CA hexameric assembly and, through two sequential oxidation steps, crosslinks two CA proteins as a molecular hinge (Figure 4A), thereby driving aberrant CA multimerization and degradation. Consistent with this mechanism, TEM and TZM-bl assays showed that ACAi-001 also exerts direct anti-HIV-1 activity against intact viral particles in vitro.

Crews *et al*. reported interesting compounds that recruit target proteins for ubiquitination and removal by the proteasome (27). Although this approach appears to be attractive for causing target protein degradation, it differs from the mechanism of ACAi-001-induced CA degradation. Misumi *et al*. (28) reported that the uncoating process requires an interaction between CA and peptidyl-prolyl isomerase (Pin1), which specifically recognizes phosphorylated Ser16 and Pro17 (29) overlapping with the ACAi-pocket. Pin1 catalyzes the isomerization of the phosphorylated Ser/Thr-Pro peptide bond (from cis to trans conformation) by binding to the phosphorylated Ser/Thr-pro peptide, initiating the uncoating process in the early-stage of HIV-1 infection. Thus, covalent binding of ACAi-001 to Ser16 may also destabilize CA proteins, such as by initiating the uncoating process; however, this requires further research.

In contrast, Eb binds covalently to the Cys-pocket (14) (Figures 2A, 5A, S5C, and S5D), which did not induce CA multimerization and degradation (Figure S3A). Collectively, these results indicate that compounds that only covalently bind to the Cys-pocket do not induce CA multimerization and degradation. When TZM-bl cells were simultaneously treated with ACAi-001 and HIV-1_NL4-3_, ACAi-001 did not significantly inhibit the early-stage of the HIV-1 life cycle. ACAi-001-induced CA multimerization required 3 h of incubation, while degradation occurred by 24 h, both in a time- and dose-dependent manner. This suggests that ACAi-001 does not exert anti-HIV-1 activity until CA multimerization occurs, despite binding to both the ACAi-pocket and Cys-pocket. Although docking simulations identified the ACAi-pocket as the primary binding site, ACAi-001 also binds Cys198 and Cys218 in the Cys-pocket, indicating limited structural selectivity (Figure 5). Unlike ACAi-001, however, Eb does not additionally target the ACAi-pocket, a difference in binding profile that likely underlies their distinct anti-HIV-1 mechanisms.

In the TZM-bl assay, ACAi-001 required an incubation period of at least 3 h to inhibit HIV-1 infection through viral particle inactivation (Figure 7H). This timeframe aligns with the onset of aberrant CA multimerization shown in Figure 2C, suggesting that the covalent cross-linking of adjacent CA monomers in the CA hexameric state by ACAi-001 is a key mechanism underlying its anti-HIV-1 activity. As a covalent inhibitor, this irreversible binding to CA may also confer the advantages of high potency and prolonged efficacy. Nevertheless, the *in vivo* toxicity of ACAi-001 warrants further investigation. Although cytotoxicity was observed only above the CC_50_ (45 µM for MT-2 and MT-4), and no effect on HSA/BSA multimerization or degradation was detected (Table 3 and Figure S2F), covalent binding to cysteine residues carries an inherent risk of non-specific, off-target reactions. The HIV-1 CA-encoding gene is relatively well conserved across HIV-1 subtypes and HIV-2 (30), consistent with our observation that ACAi-001 induced significant CA degradation in clinically isolated HIV-1 subtypes (Figure S3B) and in HIV-2_ROD_. This suggests that CA^WT^-targeting agents such as ACAi-001 may suppress replication across a broad spectrum of HIV-1 and HIV-2 strains. Notably, although reverse transcription is closely coupled with the CA uncoating process (36), ACAi-001 did not affect HIV-1 RT activity (Figure 6C), further supporting a mechanism distinct from conventional RT-dependent uncoating pathways.

This study has several limitations. First, ACAi-001-resistant HIV-1 generated through selection assays did not carry specific mutations in the Gag (CA) region (Table S1). Second, the aberrant CA multimerization and degradation induced by ACAi-001 were observed at concentrations exceeding the EC□□ (≥20 µM). Third, our docking simulations revealed distinct binding poses of ACAi-001 when bound to CA monomers versus CA dimers within hexameric assemblies (Figures 1A and 7A); the hexamer-specific configuration appears more functionally relevant and should guide future structure-activity relationship studies. Fourth, co-crystallization of ACAi-001 with CA proved technically challenging owing to the CA-degrading activity, and alternative structural approaches such as cryo-EM may be required to fully elucidate the binding mode.

## Conclusion

Taken together, these findings establish ACAi-001 as a covalent HIV-1 CA inhibitor that engages Ser16 and Cys198/Cys218 within the CA-NTD and -CTD, respectively, thereby inducing aberrant CA multimerization and degradation and disrupting the conical HIV-1 core. This dual mechanism, both covalent target engagement and direct disruption of the HIV-1 core, is mechanistically distinct from that of existing CA-targeting antiretrovirals such as PF74 and lenacapavir, which do not degrade CA. Because ACAi-001 engages CA regions that are highly conserved across HIV-1 subtypes, it may retain activity against viral strains resistant to current antiretrovirals. Overall, ACAi-001, with these unique mechanisms of action, may be a promising lead for next-generation CA-targeted therapeutics, warranting further structural optimization and comprehensive toxicological evaluation before clinical translation can be considered.

## Materials and Methods

### Cell lines

MT-2 and MT-4 cells (JCRB Cell Bank, Japan) were cultured in RPMI1640 (Gibco) with fetal bovine serum, penicillin, and kanamycin. 293T, HEK293, YSCCC, Li-7, HLE, HepG2, and COS7 cells (JCRB Cell Bank) and TZM-bl cells (NIH AIDS Reagent Program) were cultured in low-glucose DMEM with L-glutamine, phenol red (Fujifilm Wako), 10% fetal bovine serum, penicillin, and kanamycin.

### HIV strains

HIV-1_LAI_ was propagated in MT-2 cells. HIV-1_NL4-3_ and VSV-G HIV-1_ΔENV_ were produced by transfecting 293T cells with pNL4-3 or pCMV-VSV-G (Addgene) plus pNL4-3_ΔENV_, respectively (13). HIV-1_97UG029_, HIV-1_92UG037_, HIV-1_97ZA003_, and HIV_Ba-L_ were obtained from the NIH AIDS Reagent Program. Virus concentration was normalized by p24 ELISA (Lumipulse G1200, Fujirebio) after solubilization with 0.5% Triton X.

### Molecular modeling

Docking simulations were performed using SeeSAR and FlexX (BioSolveIT). Binding modes of ACAi-001 to the CA monomer (PDB: 4XFX) (39) and the CA dimer interface within the hexameric lattice (PDB: 8CKV) (40) were predicted based on the hydrophobic cavity size in the CA-NTD (PDB: 3H47). A library of 8,555,483 commercially available compounds was filtered by ADME-related properties (H-bond donors/acceptors, molecular weight, Log P, rotatable bonds) to yield 6,842,684 candidates (Figure S1). Structures were visualized using UCSF ChimeraX.

### Plasmids

CA-encoding plasmids were generated by inserting a PCR-amplified CA fragment from pNL4-3 into ApaI/XhoI-digested pCMV-Myc (Takara Bio, Myc epitope removed) for cellular expression, or into pET30a (± His tag; Novagen-Merck) for *E. coli* expression, using in-fusion cloning (Clontech). Site-directed mutagenesis (PrimeSTAR Max, Takara Bio) introduced S16E, C198A/C218A, or S16E/C198A/C218A mutations into pET30a CA.

### CA preparations expressed in the cells

Lysates of COS7 cells expressing CA^WT^ were divided into aliquots containing 100 µM of ACAi-001; one was stored immediately at −80°C, and the others were incubated at 37°C for 24, 48, or 72 h. Relative %CA was calculated by ELISA (Lumipulse G1200) at each time point.

### Protein expression and purification

CA protein purification followed a previously described procedure (5, 13). CA was expressed from pET30a CA in *E. coli* Rosetta (DE3) pLysS (Novagen) induced with IPTG for 3–5 h at 37°C. Cells were sonicated, and lysates were cleared by centrifugation before and after adding 5 M NaCl. The precipitate was resuspended in CA buffer (150 mM NaCl, 5 mM βME, 50 mM Tris-HCl, pH 7.5), filtered (0.45 µm), and purified by SEC (HiLoad 16/60 Superdex 200, GE Healthcare; AKTA Prime Plus). Monomer fractions were concentrated (Amicon Ultra-10K, Merck Millipore), and protein concentration was determined by BCA assay (Thermo Fisher) before storage at −80°C.

### TZM-bl assay

The TZM-bl assay was performed as previously described (13). Cells (NIH AIDS Reagent Program) were seeded in black 96-well plates for 24 h, then treated with drug-virus mixtures (50 ng/mL virus, 25 µg/mL DEAE-dextran; 200 µL total). After 48 h, luciferase activity was measured (D-luciferin substrate) using a FluoSTAR Omega (BMG Labtech).

### RT assay

RT activity was measured using a colorimetric assay (Roche). Recombinant HIV-1 RT (4–6 ng/well) was incubated with test compounds (DMSO, EFV 2 µM, PF74 100 µM, or ACAi-001 100 µM) and a reaction mixture containing poly(A)×(dT)_15_ template/primer, DIG-dUTP, biotin-dUTP, and dTTP for 3–4 h at 37°C. Samples were transferred to streptavidin-coated plates, incubated with anti-digoxigenin-HRP antibody, and developed with ABTS substrate. Absorbance (405 nm; Versamax, Molecular Devices) was expressed as percent inhibition relative to the DMSO control.

### WB

Western blotting (WB) was performed as previously described (32). HEK293T cells transfected with pNL4-3 (Attractene, QIAGEN) were treated with compounds from 8 h post-transfection for 48 h. Viral particles were filtered (0.22 µm), purified over a 15% sucrose-PBS cushion, normalized by p24 level, and stored at −80°C. Viral lysates (for RT/IN analysis) and HTLV-1-infected MT-2 cell lysates (for Gag analysis) were prepared after 72-h treatment with ACAi-001 (100 µM). Cells were lysed in M-PER buffer with protease inhibitors (Thermo Fisher); protein and p24 levels were quantified by BCA assay and ELISA (Lumipulse G1200), respectively. For CA multimerization assays, recombinant CA hexamers (10 µM) were incubated with ACAi-001 (0.5–50 µM) or DMSO for 24 h in high-(3 M NaCl) or low-salt (150 mM NaCl) buffer; for IP-6 assays (39), CA was pre-incubated with 100 µM IP-6 for 3–6 h before adding ACAi-001 (25–100 µM) or DMSO for 24 h. Samples were resolved by SDS-PAGE (5–20%; Nacalai Tesque), transferred to nitrocellulose/PVDF membranes, and probed with antibodies against CA, Gag, IN, RT, HTLV-1 Gag/p19, HSA, BSA, and β-actin (catalog information in Supporting Information), followed by HRP-conjugated secondary antibodies and chemiluminescent detection.

### SEC

Recombinant CA proteins were analyzed by SEC (Cosmosol-120 column, Nacalai Tesque; 1.0 mL/min in PBS, pH 7.4) as previously described (13, 33). CA (40 µM) in ammonium bicarbonate buffer (pH 7.5), with or without ACAi-001, was resolved on an Agilent 1220 Infinity LC system (280 nm detection), calibrated with ovalbumin (44 kDa) and myoglobin (17 kDa) standards. Procedures were performed at 4°C.

### CA multimerization assay

CA multimerization assays were performed as previously described (13, 18). Compounds were added to CA (30 µM) in 50 mM phosphate buffer (pH 8.0, 50 mM NaCl); assembly was initiated with 50 mM phosphate buffer (pH 8.0, 5 M NaCl), and optical density (350 nm) was measured every 5 min for 2 h (FluoSTAR Omega).

### Differential scanning fluorimetry

Differential scanning fluorimetry was performed as previously described (13, 19). Recombinant CA (25 µM) was treated with compounds for 8–12 h at 25°C, then mixed with SYPRO Orange (5×; Life Technologies) and heated from 25°C to 95°C using a 7500 Fast Real-Time PCR System (Applied Biosystems). Data are expressed as the relative fluorescence intensity ratio across this range.

### Electrospray ionization-MS and LC-MS

ESI-MS was performed as previously described (13, 33). Native MS spectra of CA with ACAi-001 were acquired on an ESI-QTOF mass spectrometer (Impact II, Bruker Daltonics; 3.3 µL/min infusion). Denatured samples were analyzed by QTOF-MS coupled to LC (Ultimate 3000 HPLC, Thermo Fisher) with a Captive Spray source (dry heater 150°C, dry gas 8.0 L/min, capillary voltage 1000 V, end plate offset −500 V; 1 Hz, 100–3000 m/z). Molecular weights were determined by protein deconvolution (Data Analysis 4.4, Bruker Daltonics) and calculated using the Peptide Mass Calculator v3.2.

### Viral core analysis

HIV-1 viral core analysis by sucrose density gradient centrifugation was performed as previously described (25, 26). Virions produced in the presence of DMSO or ACAi-001 were concentrated on a 15% sucrose cushion (35,000 rpm, 30 min, 4°C; SW55 rotor, Optima X, Beckman Coulter), envelope-stripped with 0.2% Triton-X, and loaded onto 25–70% linear sucrose gradients (STE buffer) for ultracentrifugation (28,500 rpm, 16 h, 4°C). Eleven fractions were collected and analyzed by p24 ELISA and WB with anti-CA antibody.

### TEM

Culture supernatants from COS7 cells expressing HIV^WT^ were filtered (0.22 µm) and virions pelleted by ultracentrifugation (20,000 × g, 24 h). Pellets were resuspended in PBS and incubated with 100 µM ACAi-001 or DMSO for up to 72 h, then re-pelleted as above. Virions were fixed in 2% glutaraldehyde, embedded in 2% agar, post-fixed in 2% osmium tetroxide, dehydrated through graded ethanol, and embedded in epoxy resin (Tokai Electron Microscopy Inc.). Ultrathin sections (50 nm) were stained with uranyl acetate and lead citrate and imaged by TEM (JEM-1400Plus, JEOL) with a CCD camera (VELETA, Olympus).

### Drug susceptibility assay

Susceptibility of HIV-1_LAI_ and HIV-2_ROD_ to ACAi-001 and control drugs was determined as previously described (34). MT-2 cells (10^4^/mL) were infected with 100× TCID_50_ of virus in the presence or absence of compounds and cultured for 7 days; viability and cytotoxicity (CC_50_) were assessed by MTT assay. Primary HIV-1 isolate susceptibility was assessed using PHA-stimulated PBMCs (10^6^/mL, single donor per experiment) infected at 50× TCID_50_ with serial drug dilutions. Drug-resistant laboratory strains were tested in MT-4 cells (10^5^/mL, 100× TCID_50_) as previously described (35); after 7 days, p24 levels (Lumipulse G1200) were used to calculate the 50% effective concentration relative to drug-free controls.

### Compounds

ACAi-001 contains a redox-active hydroquinone moiety; assay interference was excluded using HSA/BSA controls (Figure S2F) and by confirming antiviral activity under non-cytotoxic conditions (MTT, TZM-bl). ACAi-001 (ChemBridge) and synthesized ACAi-001, ANS-1028, and ANS-1050 were confirmed to be ≥95% pure by HPLC/NMR analysis (Supporting Information). Other reagents (RAL, zidovudine, ABC, 3TC, PF74, efavirenz, and Eb) were obtained commercially (sources listed in Supporting Information) and stored in DMSO or methanol before use.

## Acknowledgments

This work was supported by the Japan Society for the Promotion of Science, KAKENHI; grant numbers 23K24379, 18K08435, 15K09574, and 23791140 (M. Amano). This work was also supported by the Translational Research Grants from the Japan Agency for Medical Research and Development [AMED]/Center for Clinical and Translational Research of Kyushu University Hospital (A105 and A171) (M. Amano). This research was also supported by the Grants from Uehara Memorial Foundation (M. Amano), Mochida Memorial Foundation for Medical and Pharmaceutical Research (M. Amano), Takahashi Industrial and Economic Research Foundation (M. Amano), Terumo Life Sciences Foundation (M. Amano), Takeda Science Foundation (M. Amano), Sumitomo Electric Group CSR Foundation (M. Amano), and SENSHIN Medical Research Foundation (M. Amano).

We thank Drs. Toshikazu Miyakawa and Hirotomo Nakata for their valuable discussions and suggestions. We also thank Drs. Pedro Miguel Salcedo-Gómez, Rui Zhao, Travis Chia, and Ms. Sachiko Otsu for their technical support. AI-assisted proofreading was used to improve the English language quality of this manuscript using Claude (Anthropic).

## Abbreviations Used

CA: capsid
ACAi-001: N-(2,4-difluorophenyl)-2-(2,5-dihydroxybenzoyl)hydrazine-1-carbothioamide
LEN: lenacapavir
Eb: ebselen
SEC: size-exclusion chromatography
DSF: differential scanning fluorimetry
ESI-MS: electrospray ionization mass spectrometry
LC-MS: liquid chromatography-mass spectrometry
TEM: transmission electron microscopy
MTT: 3-(4,5-dimethylthiazol-2-yl)-2,5-diphenyltetrazolium bromide
ELISA: enzyme-linked immunosorbent assay
WB: Western blotting
βME: β-mercaptoethanol
IP-6: inositol hexaphosphate
ADME: absorption, distribution, metabolism, and excretion
TCID□□: 50% tissue culture infectious dose

## Conflict of Interest

Declaration of Competing Interest: The authors declare no competing financial interest.

## Data Availability

All data supporting the findings of this study are available within the article and its Supporting Information.

## Author contributions

Conceptualization: Tomofumi Nakamura, Masayuki Amano

Data curation: Tomofumi Nakamura, Nobutoki Takamune, Masaharu Sugiura, Masayuki Amano

Formal analysis: Tomofumi Nakamura, Nobutoki Takamune, Masaharu Sugiura, Masayuki Amano

## Funding acquisition: Masayuki Amano

Investigation: Tomofumi Nakamura, Nobutoki Takamune, Mayu Okumura, Masayuki Amano

Methodology: Tomofumi Nakamura, Nobutoki Takamune, Masaharu Sugiura, Masayuki Amano

Supervision: Jun-ichirou Yasunaga

Validation: Tomofumi Nakamura, Masayuki Amano

Writing – original draft: Tomofumi Nakamura

Writing – review & editing: Masayuki Amano, Masaharu Sugiura.

## Supporting Information

Figures S1–S7 showing additional experimental data including *in silico* screening procedure, CA purification and MS characterization, CA multimerization and degradation assays, βME inhibition, mutant CA analysis, TZM-bl inhibition, and anti-HIV-1 effects on viral particles; Tables S1–S2 showing Gag sequence analysis of ACAi-001-resistant HIV-1 and antiviral activity of ACAi-001 derivatives (PDF).

## Notes

### Competing Interest Statement

The authors have declared no competing interest.

### Summary of Updates

Updated the Graphical Abstract and Figures 1, 4, 5, 6, and 7 with corresponding revisions to the manuscript text.

