## Supplemental file for "A covalent anti-HIV compound induces HIV-1 capsid multimerization and degradation, decomposing the viral core"

**Author(s):**

Tomofumi Nakamura (Kumamoto University)

Nobutoki Takamune (Kumamoto University)

Mayu Okumura (Kumamoto University)

Masaharu Sugiura (Sojo University)

Jun-ichirou Yasunaga (Kumamoto University)

Masayuki Amano (Kumamoto University and Kumamoto Daiich Hospital)

**Corresponding Author:**

Tomofumi Nakamura, (T.N.)

Masayuki Amano, (M.A.)

**This file includes** Figures S1 to S7, Table S1 to S2**,** and supplemental method.

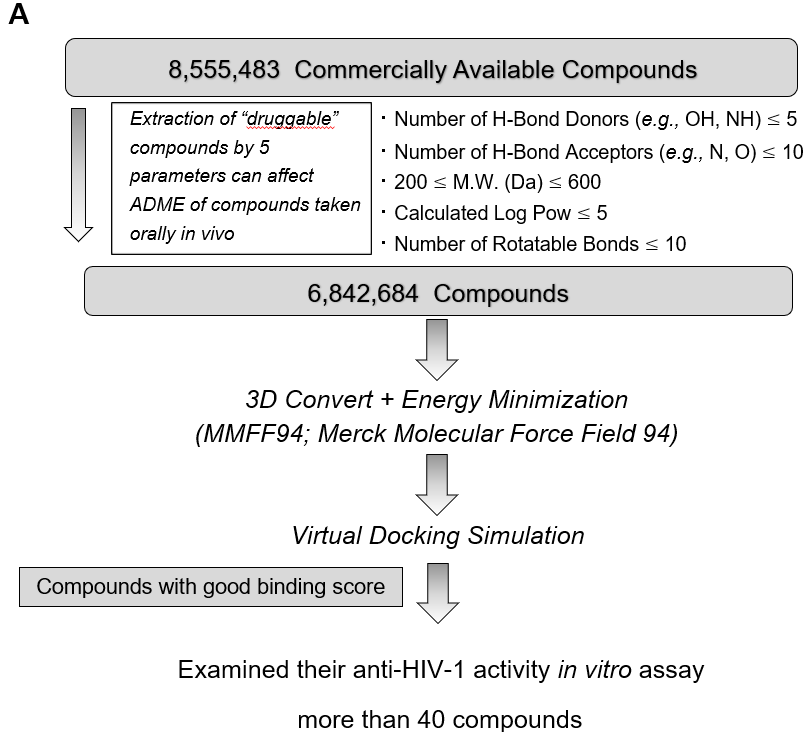

**Fig.S1 *In silico* screening procedure for ACAi-001 target cavity.** (**A**) The hydrophobic cavity was a proper size in the CA-NTD using the crystallographic data of the CA monomer and had enough space for certain small compounds to potentially fit snugly. The 6,842,684 compounds were extracted from a commercially available library of 8,555,483 compounds, which were expected to have good in vivo pharmacokinetics by the following factors; number of H-bond donors and acceptors, molecular weight, calculated Log P, and number of rotatable bonds, since these factors are related to the ADME of compounds that may be taken orally. The more than 40 compounds were evaluated for anti-HIV-1 activity using MTT in vitro assay. Docking simulations were performed using SeeSAR and FlexX version 10 (BioSolveIT GmbH, Sankt Augustin, Germany)

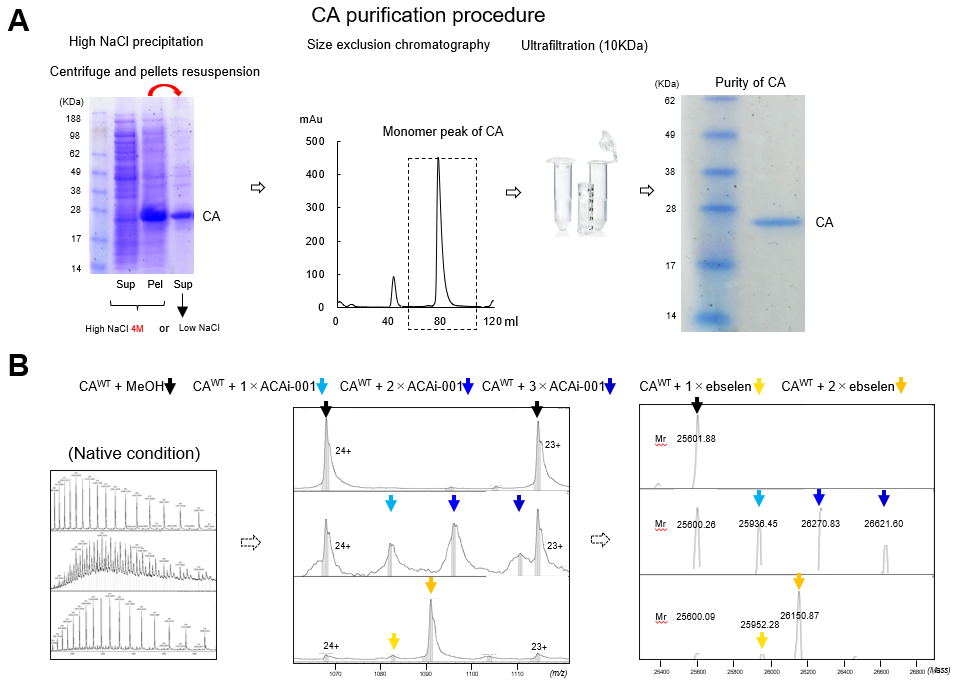

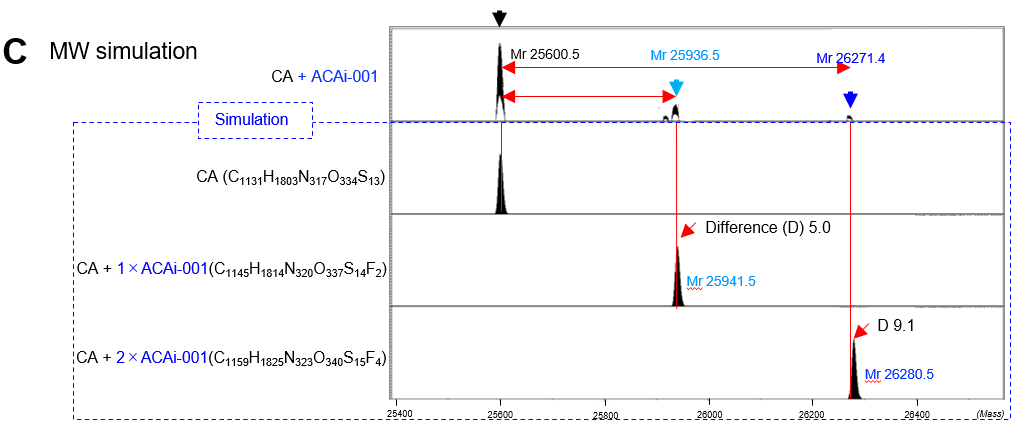

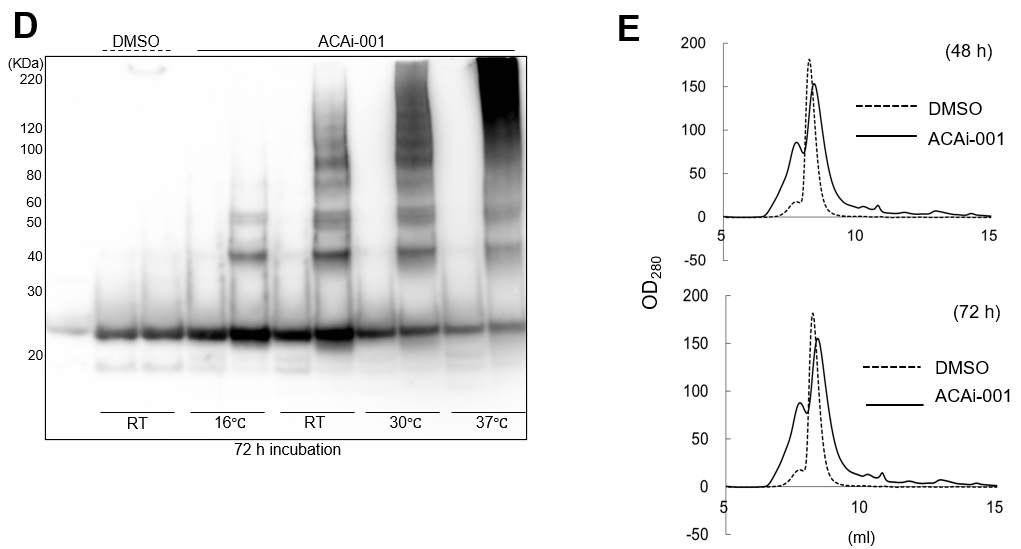

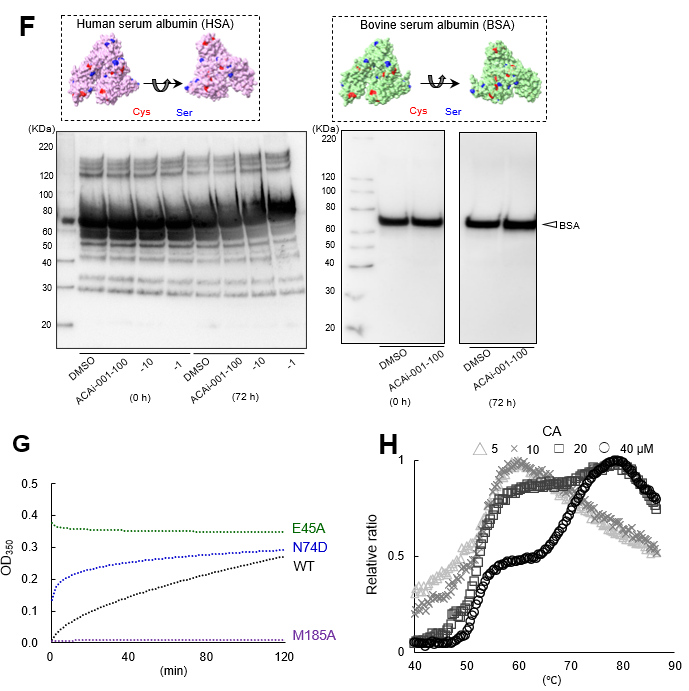

**Fig. S2 Profile of purified CA proteins induced by ACAi-001.** (**A**) Purification procedure of CA proteins expressed in *E. coli* using high sodium precipitation, SEC, and ultrafiltration. (**B**) Binding of ACAi-001 and Eb to CA in native condition using electrospray ionization-mass spectrometry (ESI-MS). (**C**) Comparison between detected and simulated MW of CA plus ACAi-001. Detected liquid chromatography-mass spectrometry (LC-MS) spectra of CA monomer covalently bound to one or two ACAi-001 are shown in the top panel. The simulated MWs of CA plus one or two ACAi-001 are shown. Red arrows indicate the differences underlying the covalent bonds between the detected and simulated MWs (D5 for one ACAi-001 bond and D9.1 for two ACAi-001 bonds). (**D**) Temperature (16°C, room temperature; RT, 30°C, and 37°C) dependent aberrant CA multimerization and degradation induced by ACAi-001 using WB compared to DMSO control. (**E**) SEC analysis of CA proteins. CA peaks treated with ACAi-001 (100 µM) for 48 and 72 h of incubation are shown (**F**) Human serum albumin (HSA) and bovine serum albumin (BSA) treated with DMSO and ACAi-001 (100 µM) for 72 h at 37°C. (**G**) Multimerization ability of purified CA carrying E45A, N74D, and M185A mutations using CA multimerization assay. (**H**) Thermal stability of CA at different concentration (5, 10, 20, and 40 µM) using differential scanning fluorimetry (DSF).

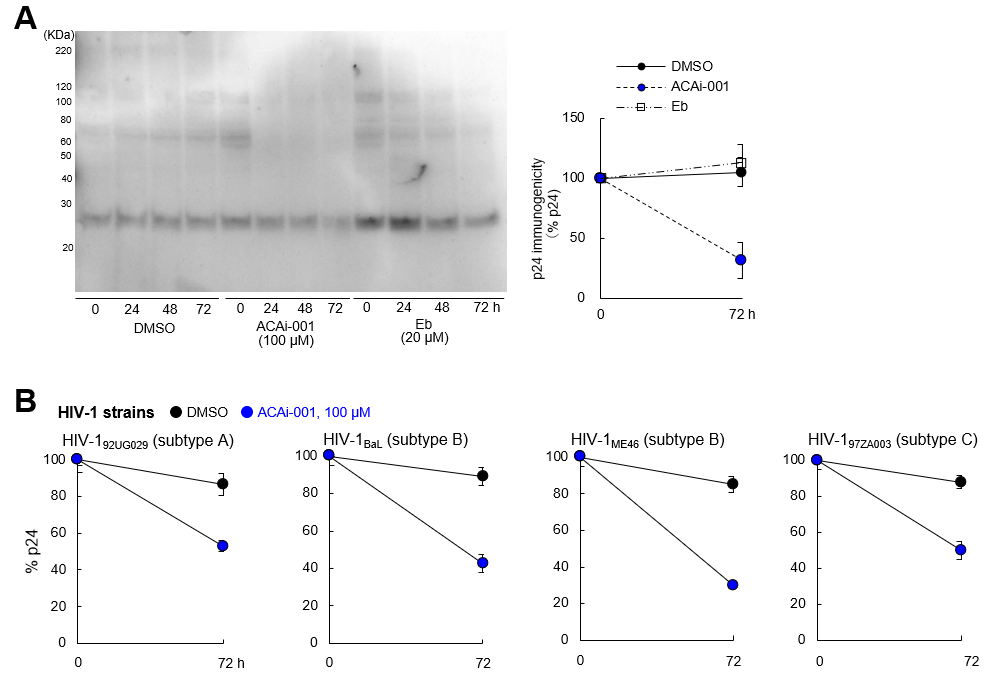

**Fig. S3 CA multimerization and degradation induced by ACAi-001 are detected by WB and p24 enzyme-linked immunosorbent assay.** (**A**) Time-dependent cell expressing CA states in the presence of DMSO (1%; black circle), ACAi-001 (100 µM; white circle), ebselen (Eb) (20 µM; black square) by the CA immunogenicity using WB and an automated ELISA device (Lumipluse G1200). (**B**) CA immunogenicity of HIV-1 clinical strains (92UG029 and BaL; subtype A, ME46; subtype B, and 97ZA003; subtype C) in the presence of DMSO (black) and ACAi-001; 100 µM (blue) incubation for 72 h at 37°C. Two or three independent experiments were performed and error bars indicate ±SD, and the representative data are shown.

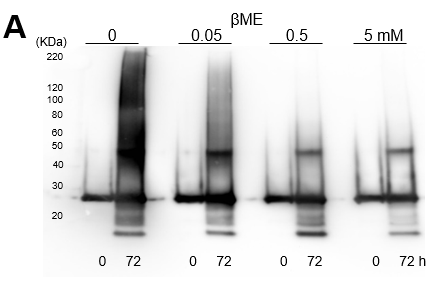

**Fig. S4 β-mercaptoethanol (βME) inhibits CA multimerization and degradation induced by ACAi-001.** (**A**) A mixture of CA proteins and ACAi-001 (100 µM) was incubated for 72 h at 37°C after the addition of different concentrations of βME (0, 0.05, 0.5, and 5 mM). Dose-dependent inhibition of βME to CA multimerization and degradation induced by ACAi-001 is shown.

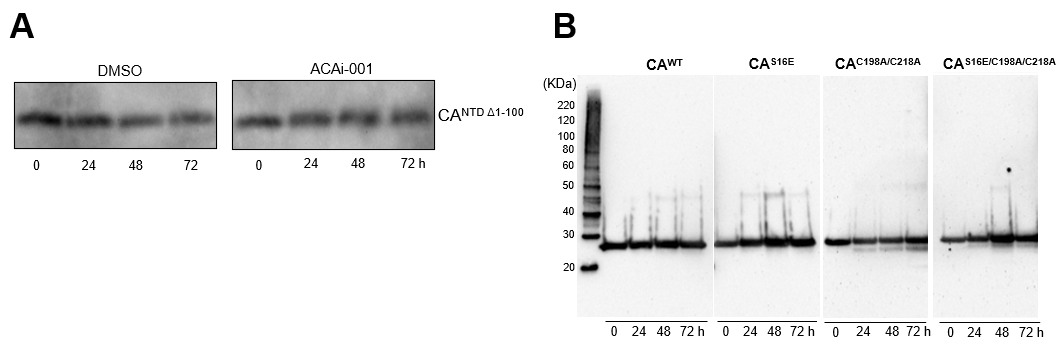

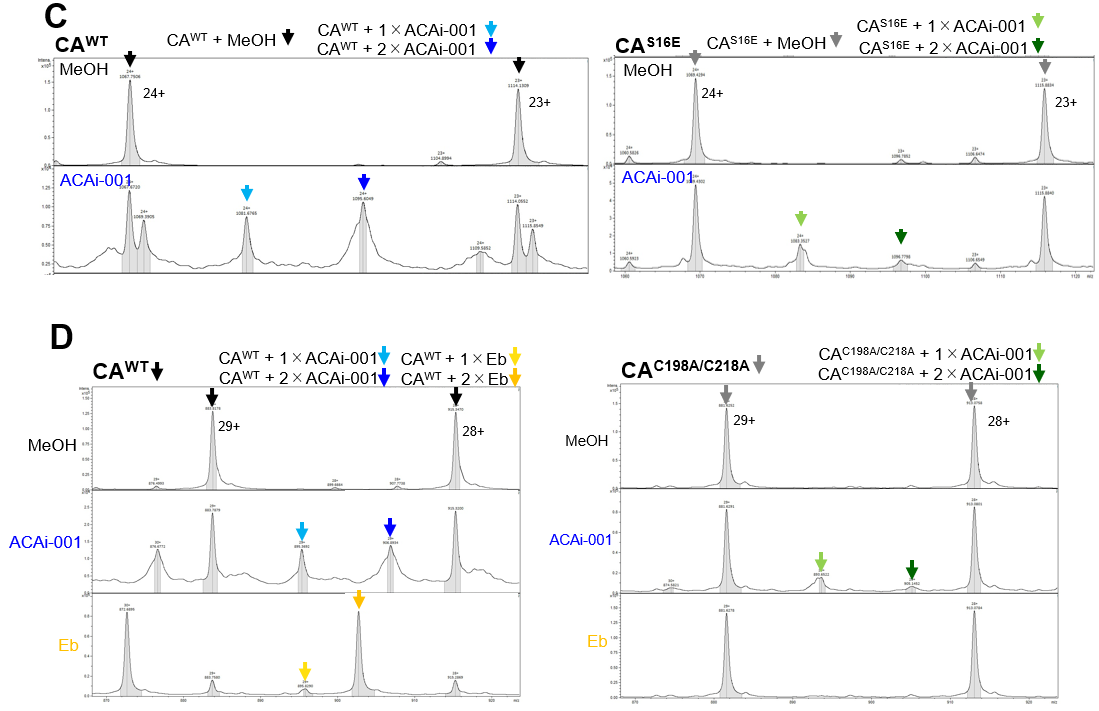

**Fig. S5 Effect of ACAi-001 on CA^NTDΔ1-100^ and covalent bonds of ACAi-001 to recombinant CA^S16E^ and CA^C198A/C218A^.** (**A**) Cell lysate containing CA^NTDΔ1-100^, deleted from the first to 100 amino acid residues of CA, was incubated for 24 h at 37°C in the presence of DMSO and ACAi-001 (100 μM), and were confirmed by WB using an anti-CA antibody. (**B**) Stability of purified CA^WT^, CA^S16E^, CA^C198A/C218A^, and CA^S16E/C198A/C218A^ during incubation at 37°C for up to 72 h (**C and D**) The binding of ACAi-001 (50 μM; light green and green arrows) to CA^S16E^ and that of ACAi-001 (50 μM; light green and green arrows) and ebselen (Eb) (50 μM; yellow and orange) to CA^C198A/C218A^ were investigated by LC-MS in the denature condition compared to CA^WT^ (1% MeOH; black arrows), CA^S16E^ (1% MeOH; gray arrows), or CA^C198A/C218A^ (1% MeOH; gray arrows). All assays were performed at least twice independently and the representative data are shown.

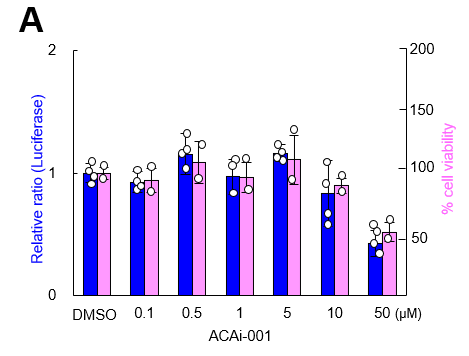

**Fig. S6 Inhibition of HIV infection and cytotoxicity to TZM-bl cells by ACAi-001.** (**A**) The inhibition of HIV-1 infectivity (early-stage inhibition) by ACAi-001, as measured by TZM-bl assay (simultaneous addition of virus and ACAi-001), is shown as the blue graphs, and the cytotoxicity of ACAi-001 to TZM-bl cells, as measured by MTT assay, is shown as the pink graphs.

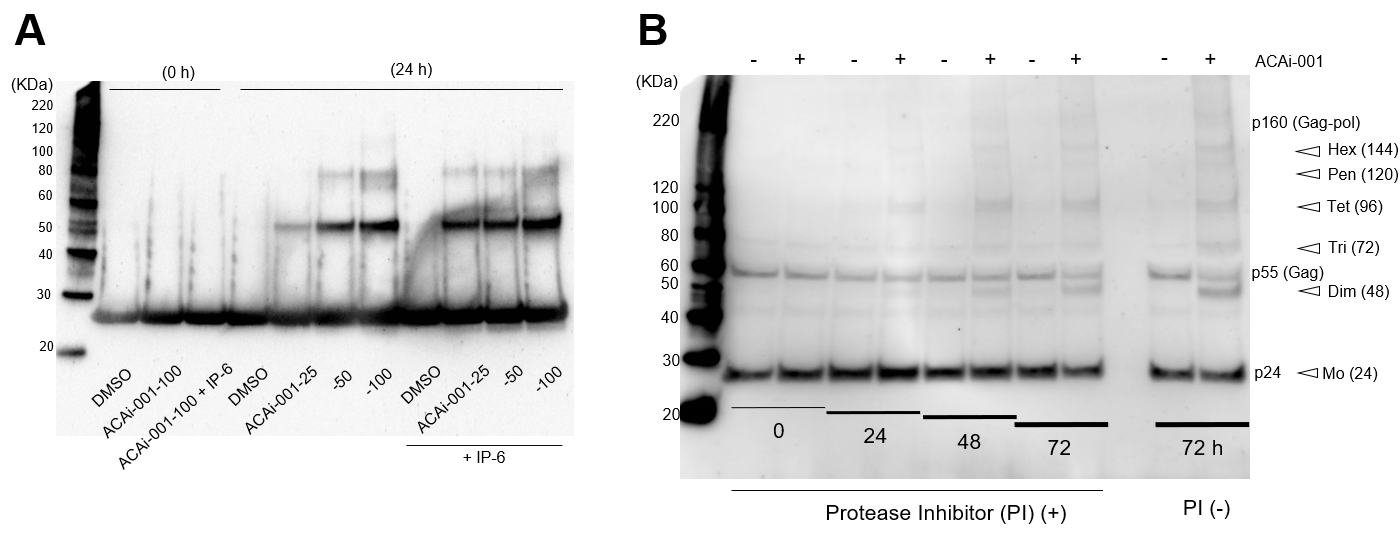

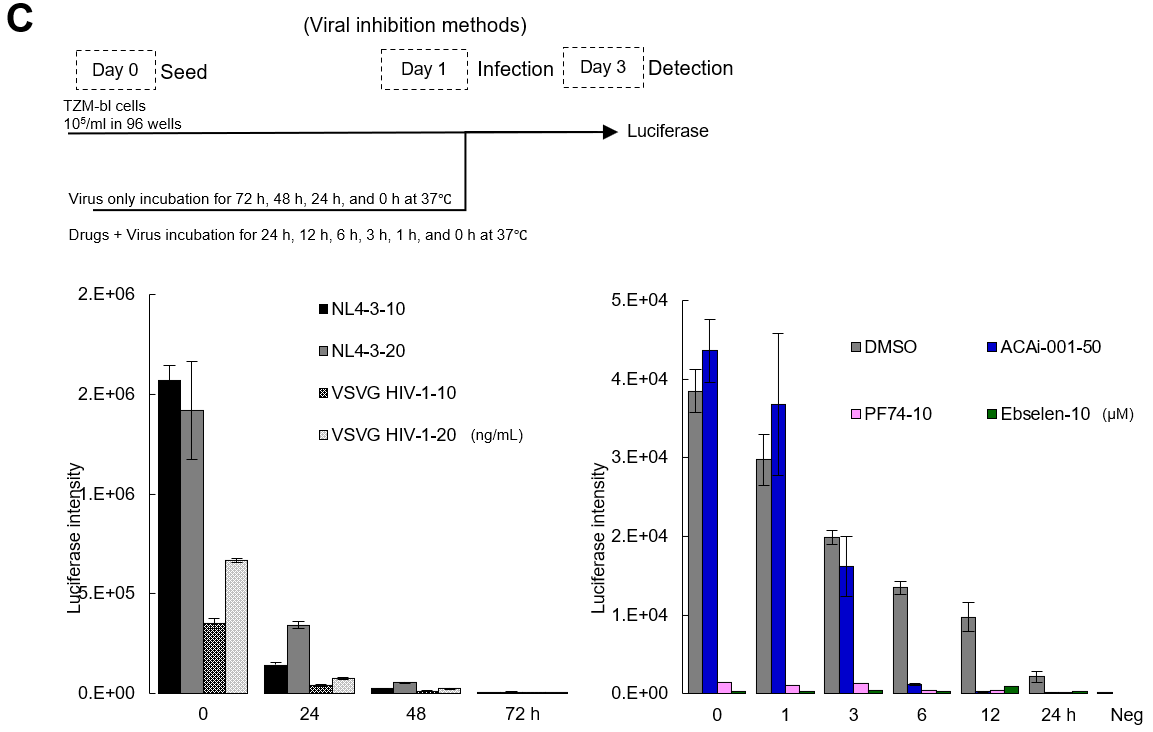

**Fig. S7Anti-HIV-1 effects of ACAi-001 on HIV-1 viral particles.** (**A**) CA hexamers were stabilized with IP-6 (3 to 6 h incubation at 37°C) prior to treatment with ACAi-001 at the indicated concentrations (25, 50, and 100 µM). After 24-h incubation at 37°C, ACAi-001 induced dose-dependent CA multimerization. (**B**) CA status of purified HIV in the presence of ACAi-001 incubated for 24, 48, and 72 h with or without protease inhibitors by WB using CA polyclonal antibody. Open arrows indicate MW of the CA multimers: monomer (Mo); 24KDa, dimer (Dim); 48KDa, trimer (Tri); 72KDa, tetramer (Tet); 96KDa, pentamer (Pen); 120KDa, and hexamer (Hex); 144Da, respectively. (**C**) TZM-bl assay procedure to determine the HIV infectivity incubated with the drugs for 0, 24, 48, and 72 h. The left bar graph shows the infectivity of HIV-1_NL4-3_ and VSV-G HIV-1_dENV_ (10 or 20 ng/mL) evaluated by TZM-bl assay after incubation for 0, 24, 48, and 72 h at 37°C. The right bar graph illustrates the infectivity of VSV-G HIV-1_dENV_ (20 ng/mL) treated with DMSO, ACAi-001 (50 µM), PF74 (10 µM), and Eb (10 µM) for 0, 1, 3, 6, 12, and 24 h at 37°C. All assays were duplicates, and error bars indicate ±SD from at least two or three independent experiments. Statistical significance was examined using Student’s t-test; *, P < 0.05, **, P < 0.005.

**Table S1. Amino acids sequence of gag region in ACAi-001 resistant HIV-1_NL43_ produced by selection assay**

1MA(p15)	100

WT MGARASVLSG GELDKWEKIR LRPGGKKQYK LKHIVWASRE LERFAVNPGL LETSEGCRQI LGQLQPSLQT GSEELRSLYN TIAVLYCVHQ RIDVKDTKEA

1 ---------- ********** ********** ********** ********** ********** ********** ******L*** ********** **********

2 *D******** ********** ********** ********** ********** ********** ********** ********** ********** **********

3 ********** ********** ********** ********** ********** ********** ********** ********** ********** **********

4 ********** ********** ********** ********** ********** ********** ********** ********** ********** **********

5 -********* ********** ********** ********** ********** ********** ********** ********** ********** **********

6 ---------- ********** ********** ********** ********** ********** ********** ********** ********** **********

7 --------** ********** ********** ********** ********** ********** ********** ********** ********** ******A***

8 ---------- -----***** ********** ********** ********** ********** ********** ********** ********** **********

9 ---------- -********* ********** ********** ********** ********** ********** ********** ********** **********

10 -********* ********** ********** ********** ********** ********** ********** ********** ********** **********

101	CA(p24)	200

LDKIEEEQNK SKKKAQQAAA DTGNNSQVSQ NYPIVQNLQG QMVHQAISPR TLNAWVKVVE EKAFSPEVIP MFSALSEGAT PQDLNTMLNT VGGHQAAMQM

1 ********** ********** ********** ********** ********** ********** ********** ********** ********** **********

2 ********** ********** ********** ********** ********** ********** ********** ********** ********** **********

3 ********** ********** ********** ********** ********** ********** ********** ********** ********** **********

4 ********** ********** ********** ********** ********** ********** ********** ********** ********** **********

5 ********** ********** ********** ********** ********** ********** ********** ******K*** ********** **********

6 ********** ********** ********** ********** ********** ********** ********** ********** ********** **********

7 ********** ********** ********** ********** ********** ********** ********** ********** ********** **********

8 ********** ********** ********** ********** ********** ********** ********** ********** ********** **********

9 ********** ********** ********** ********** ********** ********** ********** ********** ********** **********

10 ********** ********** ********** ********** ********** ********** ********** ********** ********** **********

201	300

LKETINEEAA EWDRLHPVHA GPIAPGQMRE PRGSDIAGTT STLQEQIGWM THNPPIPVGE IYKRWIILGL NKIVRMYSPT SILDIRQGPK EPFRDYVDRF

1 ********** ********** ********** ********** ********** ********** ********** ********** ********** **********

2 ********** ********** ********** ********** ********** ********** ********** ********** ********** **********

3 ********** ********** ********** ********** ********** ********** ********** ********** ********** **********

4 ********** ********** ********** ********** ********** ********** ********** ********** ********** **********

5 ********** ********** ********** ********** ********** ********** ********** ********** ********** **********

6 ********** ********** ********** ********** ********** ********** ********** ********** ********** **********

7 ********** ********** ********** ********** ********** ********** ********** ********** ********** **********

8 ********** ********** ********** ********** ********** ********** ********** ********** ********** **********

9 ********** ********** ********** ********** ********** ********** ********** ********** ********** **********

10 ********** ********** ********** ********** ********** ********** ********** ********** ********** **********

301	NC(p7)	400

YKTLRAEQAS QEVKNWMTET LLVQNANPDC KTILKALGPG ATLEEMMTAC QGVGGPGHKA RVLAEAMSQV TNPATIMIQK GNFRNQRKTV KCFNCGKEGH

1 ********** ********** ********** ********** ********** ********** ********** ********** ********** **********

2 ********** ********** ********** ********** ********** ********** ********** ********** ********** **********

3 ********** ********** ********** ********** ********** ********** ********** ********** ********** **********

4 ********** ********** ********** ********** ******I*** ********** ********** ********** ********** **********

5 ********** ********** ********** ********** ********** ********** ********** ********** ********** **********

6 ********** ********** ********** ********** ********** ********** ********** ********** ********** **********

7 ********** ********** ********** ********** ********** ********** ********** ********** ********** **********

8 ********** ********** ********** ********** ********** ********** ****K*I*** ******I*** ******K*** **********

9 ********** ********** ********** ********** ********** ********** ********** ********** ********** **********

10 ********** ********** ********** ********** ********** ********** ********** ********** ********** **********

401	p1	p6	500

IAKNCRAPRK KGCWKCGKEG HQMKDCTERQ ANFLGKIWPS HKGRPGNFLQ SRPEPTAPPE ESFRFGEETT TPSQKQEPID KELYPLASLR SLFGSDPSSQ

1 ********** ********** ********** ********** ********** ********** ********** ********** ********** **********

2 ********** ********** ********** ********** ********** ********** ********** ********** ********** **********

3 ********** ********** ********** ********** ********** ********** ********** ********** ********** **********

4 ********** ********** ********** ********** ********** ********** ********** ********** ********** **********

5 ********** ********** ********** ********** ********** ********** ********** ********** ********** **********

6 ********** ********** ********** ********** ********** ********** ********** ********** ********** **********

7 ********** ********** ********** ********** ********** ********** ********** ********** ********** **********

8 ********** ********** ********** ********** ********** ********** ********** ********** ********** **********

9 ********** ********** ********** ********** ********** *K*K****** ********** ********** ********** **********

10 ********** ********** ********** ********** ********** ********** ********** ********** ********** **********

A total of ten sequences of Gag (MA, CA, p2, NC, p1, and p6) derived from the ACAi-001-resistant HIV-1 (replicable HIV-1 in 50 µM ACAi-001) were identified, and AA mutations are shown in red compared to the wild-type HIV-1_NL4-3_. *, wild type AA; -, not determined.

**Table S2. Anti-HIV-1 activity of ACAi-028, ANS-1028, and ANS-1050 against HIV-1 and HIV-2**

| Virus | Cells | EC_50_ (μM) | | | |
| --- | --- | --- | --- | --- | --- |
|  |  | ACAi-028 | ANS-1028 | ANS-1050 | Ebselen |
| HIV-1_LAI_ | MT-2 | 0.55 ± 0.04 | >10 | >10 | 1.73 ± 0.04 |
| HIV-2_ROD_ | MT-2 | >10 | >10 | >10 | 8.16 ± 0.46 |

MT-2 cells (10^4^/ml) were exposed to 100 TCID_50_ of HIV-1_LAI_ or HIV-2_ROD_ and cultured in the presence of various concentrations of each compound, and the EC_50_ (50% effective concentration) values were determined using the MTT assay.

**Supporting information methods**

**Generation and analysis of drug-resistant HIV-1 variants**.

Selection experiments for resistant HIV-1 variants were performed. In brief, MT-4 cells (1.0×10^5^ cells/ml) were exposed to HIV-1_NL4-3_ (50% tissue culture infective doses [TCID_50_s]) and cultured in the presence of ACAi-001, at an initial concentration of the EC_50_. Viral replication in MT-4 cells was monitored by p24 levels in the culture medium at intervals of 1 week. The selection procedure was continued until ACAi-001 concentration was over 50 µM. Proviral DNA sequences of the ACAi-001 resistant HIV-1 were extracted from the infected MT-4 cells using a NucleoSpin tissue kit (TaKaRa, Japan). DNA sequences of the Gag regions were amplified using the following primers (MAF, 5′-ATG GGT GCG AGA GCG TCG GTA TTA AGC-3′ and P6R, 5′-TTA TTG TGA CGA GGG GTC GCT GCC-3′). The PCR Gag fragment was inserted into T-Vector pMD20s (TAKARA, Japan), and the cloned T-vectors carrying the Gag sequence were analyzed using the following sequence primers (CAF1, 5′-AGG GCC TAT TGC ACC AGG CCA G -3′ and CAR1, 5′- AAG CTT GCT CGG CTC TTA GAG TTT TAT AG-3′).

**Antibodies and reagents.**

Primary antibodies used for WB were as follows: anti-CA (monoclonal, ab63913, Abcam, Cambridge, UK; polyclonal, Cat# 13-203-000, Advanced Biotechnologies, Columbia, MD, USA), anti-HIV-1 Gag (ab63917, Abcam), anti-HIV-1 IN (ab66645, Abcam), anti-HIV-1 RT (65-001, Bio Academia Co., Ltd., Osaka, Japan), anti-HTLV-1 Gag/p19 (sc-57868, Santa Cruz Biotechnology, Dallas, TX, USA), anti-HSA (F-10) (sc-271605, Santa Cruz Biotechnology), anti-BSA (2A3E6, Santa Cruz Biotechnology), and HRP-conjugated anti-beta-actin (ab49900, Abcam). HRP-conjugated anti-mouse and anti-rabbit secondary antibodies were from MBL Co., Ltd. (Tokyo, Japan). Control and reference compounds were obtained as follows: raltegravir (RAL) from Selleck Chemicals (Houston, TX, USA); zidovudine, abacavir (ABC), lamivudine (3TC), PF74, and efavirenz (EFV) from Sigma-Aldrich (St. Louis, MO, USA); and ebselen (Eb) from AdipoGen Life Sciences (Füllinsdorf, Switzerland).

**Preparation of ACAi-001.**

To a solution of 2,4-difluorophenyl isothiocyanate (79.0 mg, 0.462 mmol) in dry dichloromethane (1 mL) and dry THF (2 mL) was added 2,5-dihydroxybenzohydrazide (66.7 mg, 0.397 mmol) at rt. The mixture was stirred at rt for 4 h. Silica gel (300 mg) was added and the mixture was concentrated under reduced pressure. The residue was purified by column chromatography on silica gel (hexane/AcOEt = 1:2). The fractions containing the desired product were collected, concentrated, and filtered with diethyl ether to give ACAi-001 (73.5 mg, 0.217 mmol, 54.7% yield).

***N*-(2,4-Difluorophenyl)-2-(2,5-dihydroxybenzoyl)hydrazine-1-carbothioamide (ACAi-001).**

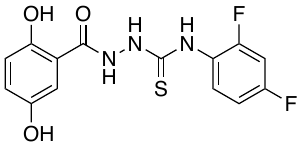
TLC: R*_f_* 0.40 (hexane/AcOEt = 1:2, stained blue with phosphomolybdic acid). ^1^H NMR (500 MHz, DMSO-d_6_, 80 °C) d 10.81 (brs, 1H), 10.70 (brs, 1H), 10.02 (brs, 1H), 9.42 (brs, 1H), 8.81 (brs, 1H), 7.56 (brs, 1H), 7.31 (d, *J* = 2.9 Hz, 1H), 7.21 (ddd, *J* = 10.3, 9.2, 2.9 Hz, 1H), 7.04 (dddd, *J* = 8.6, 8.6, 2.9, 1.2 Hz, 1H), 6.88 (dd, *J* = 8.6, 2.9 Hz, 1H), 6.79 (d, *J* = 8.6 Hz, 1H).

**Preparation of ANS-1028.**

To a solution of 2,4-difluorophenyl isothiocyanate (41.0 mg, 0.240 mmol) in dry THF (2 mL) was added 3-hydroxybenzohydrazide (30.1 mg, 0.198 mmol) at rt. The mixture was stirred at rt for 4 h and concentrated under reduced pressure. The crystalline residue was collected by filtration with hexane/AcOEt (1:1) to give ANS-1028 (61.2 mg, 0.189 mmol, 95.5% yield).

***N*-(2,4-Difluorophenyl)-2-(2,5-dihydroxybenzoyl)hydrazine-1-carbothioamide (ANS-1028).**

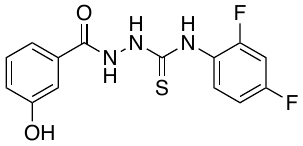
TLC: R*_f_* 0.34 (hexane/AcOEt = 1:2, stained blue with phosphomolybdic acid). ^1^H NMR (500 MHz, DMSO-d_6_, 80 °C) d 10.28 (brs, 1H), 9.66 (brs, 1H), 9.44 (brs, 1H), 9.35 (brs, 1H), 7.48 (brs, 1H), 7.38 (d, *J* = 7.7 Hz, 1H), 7.35 (dd, *J* = 2.3, 1.7 Hz, 1H), 7.27 (t, *J* = 7.7 Hz, 1H), 7.19 (ddd, *J* = 10.3, 9.2, 2.9 Hz, 1H), 7.04 (dddd, *J* = 8.6, 8.6, 2.9, 1.2 Hz, 1H), 6.97 (dd, *J* = 8.0, 2.3 Hz, 1H).

**Preparation of ANS-1050.**

To a solution of 2,4-difluorophenyl isothiocyanate (28 µL, 0.22 mmol) in dry THF (2 mL) was added 2-hydroxybenzohydrazide (31.0 mg, 0.204 mmol) at rt. The mixture was stirred at rt for 4 h and concentrated under reduced pressure to half its volume. The residue was purified by column chromatography on silica gel (hexane/AcOEt = 1:2). The fractions containing the desired product were collected, concentrated, and filtered with hexane/AcOEt (1:1) to give ANS-1050 (51.5 mg, 0.159 mmol, 77.9% yield).

***N*-(2,4-Difluorophenyl)-2-(2-hydroxybenzoyl)hydrazine-1-carbothioamide (ANS-1050).**

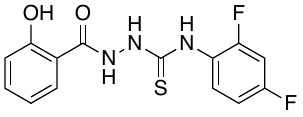
TLC: R*_f_* 0.40 (hexane/AcOEt = 1:2, stained blue with phosphomolybdic acid). ^1^H NMR (500 MHz, DMSO-d_6_, 80 °C) d 11.51 (brs, 1H), 10.86 (brs, 1H), 9.97 (brs, 1H), 9.46 (brs, 1H), 7.91 (d, *J* = 8.0 Hz, 1H), 7.54 (brs, 1H), 7.43 (apparent t, *J* = 7.7 Hz, 1H), 7.21 (ddd, *J* = 10.3, 9.2, 2.9 Hz, 1H), 7.05 (dddd, *J* = 8.6, 8.6, 2.9, 1.2 Hz, 1H), 6.96 (d, *J* = 8.0 Hz, 1H), 6.93 (t, *J* = 7.7 Hz, 1H).
